# Protein Deacetylase CobB Interplays with c-di-GMP

**DOI:** 10.1101/362293

**Authors:** Zhaowei Xu, Hainan Zhang, Xingrun Zhang, Chengxi Liu, Hewei Jiang, Fanlin Wu, Lili Qian, Daniel M. Czajkowsky, Shujuan Guo, Lijun Bi, Shihua Wang, Haitao Li, Minjia Tan, Lei Feng, Jingli Hou, Sheng-ce Tao

**Affiliations:** Key Laboratory of Systems Biomedicine (Ministry of Education), Shanghai Center for Systems Biomedicine, Shanghai Jiao Tong University, 800 Dongchuan Road, Shanghai 200240, China; State Key Laboratory of Oncogenes and Related Genes, Shanghai 200240, China; School of Biomedical Engineering, Shanghai Jiao Tong University, Shanghai 200240, China; MOE Key Laboratory of Protein Sciences, Center for Structural Biology, School of Life Sciences, Tsinghua University, Beijing 100084, China; Department of Basic Medical Sciences, School of Medicine, Tsinghua University, Beijing 100084, China; The Chemical Proteomics Center and State Key Laboratory of Drug Research, Shanghai Institute of Materia Medica, Chinese Academy of Sciences, Shanghai 201203, China; National Key Laboratory of Biomacromolecules, Key Laboratory of Non-Coding RNA and Key Laboratory of Protein and Peptide Pharmaceuticals, Institute of Biophysics, Chinese Academy of Sciences, Beijing 100101, China; School of Stomatology and Medicine, Foshan University, Foshan 528000, Guangdong Province, China; School of Life Science, Fujian Agriculture and Forestry University, Fuzhou, Fujian 350002, China; Instrumental Analysis Center, Shanghai Jiao Tong University, 800 Dongchuan Road, Shanghai 200240, China

**Keywords:** c-di-GMP, CobB, diguanylate cyclase, protein acetylation, negative feedback loop

## Abstract

As a ubiquitous bacterial secondary messenger, c-di-GMP plays key regulatory roles in processes such as bacterial motility and transcription regulation. CobB is the Sir2 family protein deacetylase that controls energy metabolism, chemotaxis and DNA supercoiling in many bacteria. Using an *E.coli* proteome microarray, we found that c-di-GMP strongly binds to CobB. Protein deacetylation assays showed that c-di-GMP inhibits CobB activity and thereby modulates the biogenesis of acetyl-CoA. Through mutagenesis studies, residues R8, R17 and E21 of CobB were shown to be required for c-di-GMP binding. Next, we found that CobB is an effective deacetylase of YdeH, a major diguanylate cyclase (DGC) of *E.coli* that is endogenously acetylated. Mass spectrometry analysis identified YdeH K4 as the major site of acetylation, and it could be deacetylated by CobB. Interestingly, deacetylation of YdeH enhances its stability and cyclase activity in c-di-GMP production. Thus, our work establishes a novel negative feedback loop linking c-di-GMP biogenesis and CobB-mediated protein deacetylation.

## Introduction

Cyclic diguanosine monophosphate (c-di-GMP) was first identified in *Gluconacetobacter xylinus*, where it was found to regulate cellulose synthesis^1^. Subsequently, c-di-GMP was shown to be involved in a wide range of bacterial biological processes such as bacterial motility, biofilm formation, virulence and transcription regulation^2-4^. However, these processes likely represent only a portion of the full scope of its diverse functions in the cell, owing to the typical challenges of unambiguously ascribing functionality directly to the activity of a specific small molecule second messenger^5^. A first step toward this understanding has often emerged from an identification of the proteins to which it strongly interacts, as exemplified in studies of c-di-GMP binding to YcgR^6^, CckA^7^, BldD^8^ and CheY-like (Cle) proteins^9^. Thus, we speculated that a better understanding of the full repertoire of c-di-GMP functions in bacteria would emerge from a comprehensive knowledge of the complete range of proteins to which it binds.

c-di-GMP is synthesized by diguanylate cyclases (DGCs)^10^ and degraded by specific phosphodiesterases (PDEs)^11-13^. In *E.coli*, the dominant DGCs are YdeH (also known as DgcZ)^14^ and DosC^15^, and the major PDEs are YhjH^16^ and DosP^15^. Each of these proteins has been shown to be modulated by “first” messengers such as light, oxygen and temperature^2^. Yet, owing to the involvement of c-di-GMP in the aforementioned fundamental biological processes that are all well-regulated by many basic metabolic mechanisms^17, 18^, there is also the possibility that c-di-GMP biosynthesis is also regulated via metabolism-related intracellular signals.

The Sir2 family protein CobB is a NAD^+^-dependent deacetylase that is highly conserved in prokaryotes^19, 20^. In *E.coli*, CobB is the sole Sir2 homolog, although there are two forms of CobB, namely, CobB and CobB_S_, the former of which has an additional 37 aa N-terminal tail^21^. CobB exhibits protein deacetylation activity and it regulates a variety of physiological functions. For example, CobB deacetylates lysine-609 of acetyl-coenzyme A synthetase (Acs) to activate its activity^22^, resulting in an increased cellular concentration of acetyl-coenzyme A (acetyl-CoA), which is a central component of energy metabolism. In addition, CobB regulates *E.coli* chemotaxis by deacetylating the chemotaxis response regulator protein (CheY)^23^, as well as the activity of N-hydroxyarylamine O-acetyltransferase (NhoA)^24^ and topoisomerase I (TopA)^25^. Yet, despite the critical role that CobB plays in many biological processes, its inherent regulation remains poorly understood^26-28^.

To globally identify c-di-GMP effectors and explore new functions of c-di-GMP, we employed an *E.coli* proteome microarray^29^ for proteome-wide identification of c-di-GMP binding proteins. Surprisingly, we found that c-di-GMP strongly binds to CobB. Subsequent biochemical analysis confirmed that c-di-GMP binds to CobB and inhibits its deacetylation activity both *in vitro* and *in vivo*. Furthermore, we found that the major DGC in *E.coli*, YdeH, is endogenously acetylated and CobB promotes the stability and activity of YdeH through deacetylation of lysine-4. Altogether, we have established an evidence-based regulation loop underlying the cytoplasmic concentration of c-di-GMP that involves its direct binding to, and thereby inhibition of CobB.

## Results

### Protein deacetylase CobB is a novel c-di-GMP effector

To identify novel effectors of c-di-GMP, an *E.coli* proteomic microarray was probed with biotin-c-di-GMP. In this way, CobB (specifically, the version of CobB with the extra N-terminal tail) was identified as a strong binder of c-di-GMP **(Fig. 1a)**. To validate this interaction, we developed a simple *in vitro* assay in where purified CobB was incubated with biotin-c-di-GMP, UV crosslinked, and then probed with fluorescent streptavidin.^30, 31^ We found that c-di-GMP indeed exhibits strong binding to CobB, although trace bindings were also observed for cGMP and c-di-AMP **(Fig. 1b)**. Mixing c-di-GMP with biotin-c-di-GMP before the two were added to CobB resulted in a significant reduction in biotin-c-di-GMP binding to CobB **(Fig. 1b)**. To quantify the binding strength of this interaction, we performed Isothermal Titration Calorimetry (ITC) and found that the c-di-GMP/CobB affinity constant, *K_d_*, is 21.6 μM with a binding stoichiometry of 0.97. We also performed ITC assays with cGMP and c-di-AMP, and no binding was detected with these two molecules **(Fig. 1c and supplementary Fig. 1a-c)**. To confirm these results, we also determined the affinity constant using Microscale Thermophoresis (MST), which yielded a *K_d_* of 22.7 ± 1.63 μM for the interaction between c-di-GMP and CobB **(Supplementary Fig. 2)**. We note however that while the affinity constants determined from ITC and MST are consistent with each other, we observed some polymerization of CobB in the presence of c-di-GMP **(Supplementary Fig. 3)**, which may somewhat affect the measurement of the affinity for c-di-GMP. Nonetheless, taken together, these results clearly show that c-di-GMP specifically binds to CobB.

**Figure 1.**
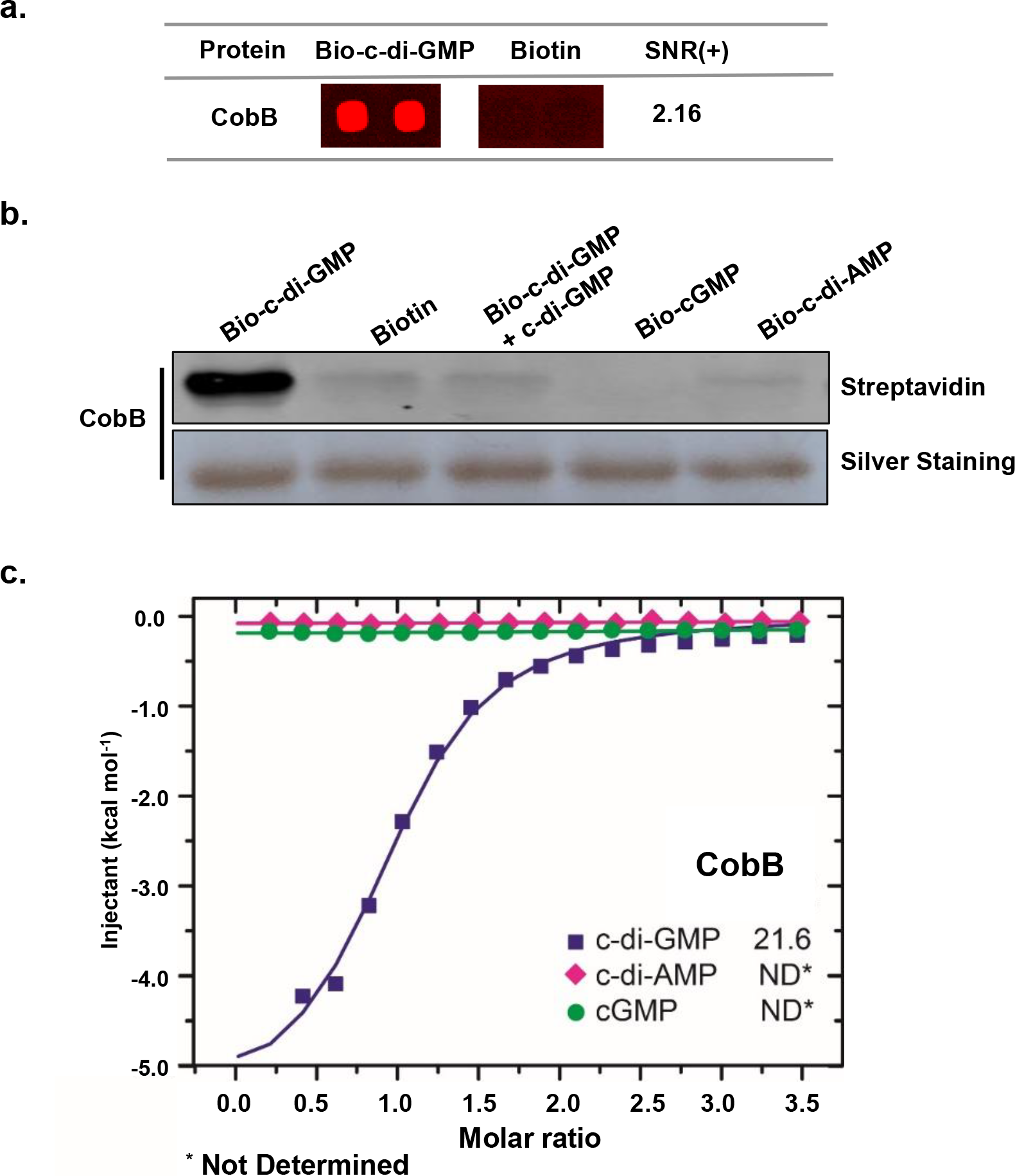
Protein deacetylase CobB is a novel c-di-GMP effector. **(a)**The *E.coli* proteome microarrays were probed with bio-c-di-GMP, followed by incubation with Cy3-conjugated streptavidin. A control experiment was carried out with biotin. Obvious binding difference of CobB on the microarrays incubated with bio-c-di-GMP vs. biotin was observed. There are two spots per protein, SNR (+) is the average signal to noise ratio of the two duplicate spots. **(b)** Streptavidin blotting assay. Affinity purified CobB was incubated with 10 μM bio-c-di-GMP. 10 μM biotin, biotin-cGMP and biotin-c-di-AMP were included as negative controls. In the 3^rd^ lane, CobB was incubated with 10 μM bio-c-di-GMP and 20 μM c-di-GMP. The bindings of the biotinylated ligands were visualized by streptavidin. Silver staining showed equal amounts of CobB were included for each reaction. **(c)** ITC analysis of the binding between c-di-GMP and CobB. 1.5 mM c-di-GMP (purple line) was titrated into CobB, in parallel assays, equal amount of cGMP (green line) and c-di-AMP (red line) were included as the negative controls. The Kd of c-di-GMP and CobB binding was determined as 21.6 μM.

### c-di-GMP inhibits the deacetylase activity of CobB and down-regulates the biogenesis of acetyl-CoA

With the demonstration of binding between c-di-GMP and CobB, we speculated that c-di-GMP may affect the deacetylase activity of CobB. To test this, we examined the activity of CobB in the presence of c-di-GMP by monitoring the deacetylation of the well-known CobB substrates, Acs^22^, CheY^23^, NhoA^24^. Before incubation with CobB, we found that purified Acs from a CobB-deficient *E.coli* (Δ*cobB*) is highly acetylated (**Fig. 2a**). Incubation with CobB resulted in significant reduction in acetylation, consistent with previous work^22^ (**Fig. 2a**). However, in the presence of c-di-GMP, the deacetylation of Acs was significantly inhibited in a dose dependent manner **(Fig. 2a and supplementary Figure 4a)**. By contrast, neither the presence of cGMP nor c-di-AMP affected the CobB deactylation of Acs. Similar results were obtained for CheY **(Supplementary Figure 4b)** and NhoA **(Supplementary Figure 4c)**.

**Figure 2.**
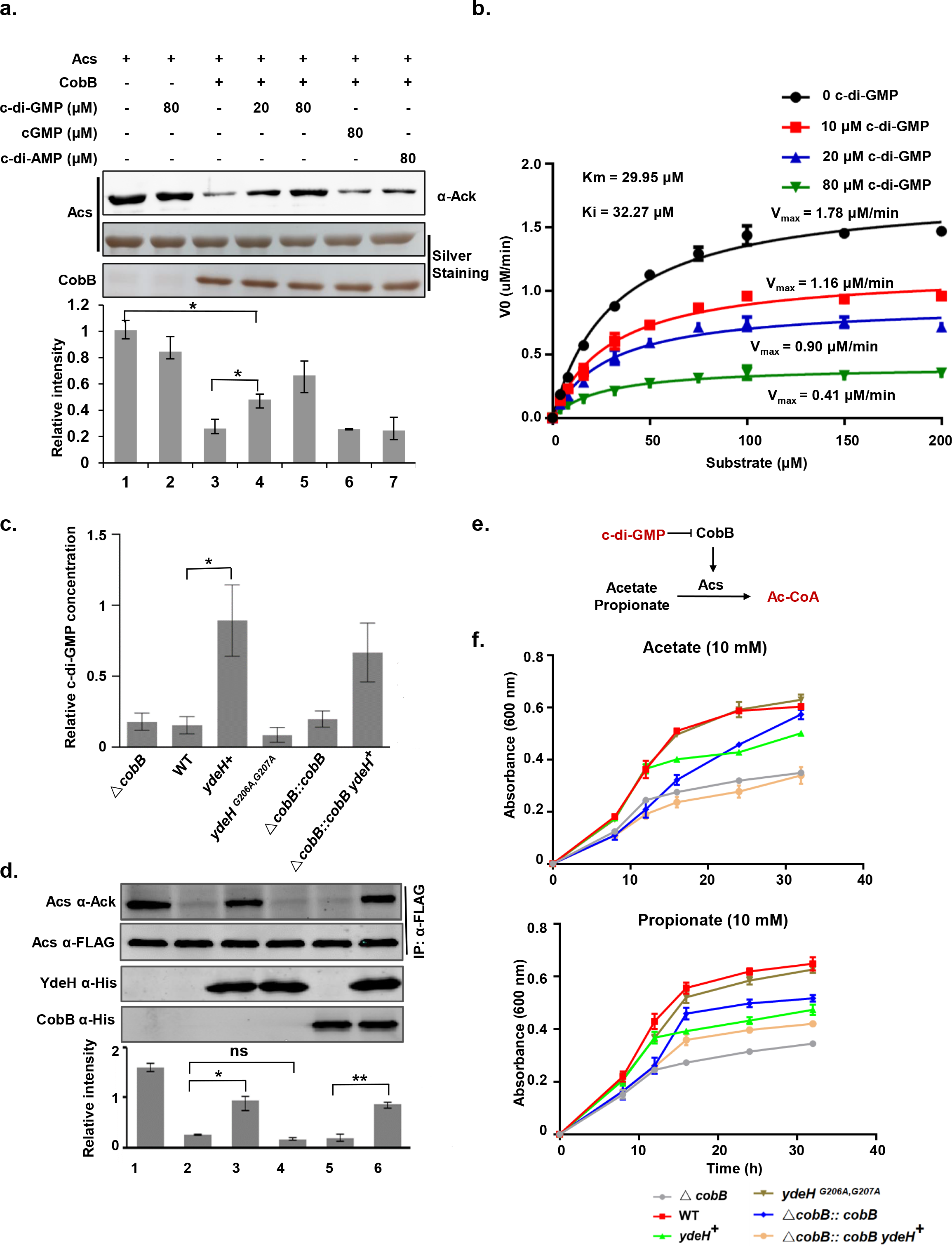
c-di-GMP inhibits the deacetylase activity of CobB and down-regulates the biogenesis of acetyl-CoA. **(a)**CobB activity assay was performed using Acs as substrate. The loss of the acetylation of Acs indicates CobB’s deacetylase activity, as shown by the pan anti-acetyl antibody. (three preparations; \*\**P* < 0.01, two-tailed Student’s t-test). The protein levels of Acs and CobB were determined by silver staining. **(b)** The kinetics of CobB enzyme-catalyzed reactions. The CobB catalytic kinetics were performed with the addition of 0, 10, 20, 80 μM c-di-GMP and an acetylated peptidewas used as the substrate. The acetylated and deacetylated peptides were quantified by HPLC with three replicates and these curves were fitting by michaelis-menten equation using GraphPad Prism 6. **(c)** The endogenous c-di-GMP levels of the 6 *E.coli* strains were determined by UPLC-IM-MS (three preparations; \**P* < 0.05, two-tailed Student’s t-test). **(d)** CobB’s deacetylation activity was monitored using endogenous Acs as substrate for the 6 strains. 3xFLAG-tagged Acs was enriched by an anti-FLAG antibody and the acetylation level was determined by the pan anti-acetyl antibody. The expression of DGC and CobB were determined using an anti-His antibody. The bar graph showed the quantitation of the acetylation level of Acs. (three preparations; \**P* 0.05 and \*\**P* < 0.01, two-tailed Student’s t-test) **(e)** c-di-GMP modulates the synthesis of acetyl-CoA through inhibiting CobB to deacetylate Acs. **(f)** The growth curves of the 6 strains were determined. The *E.coli* strains were cultured in Vogel-Bonner medium. The growth was measured at 8, 12, 16, 20, 24 and 32 h with three replicates.

To examine the inhibition of c-di-GMP on CobB in more detail, we measured the kinetics of CobB deacetylation of an acetylated peptide^32^at different concentrations of c-di-GMP. We found that c-di-GMP significantly reduces the maximal catalytic rate of CobB, with no changes to the Km values, yielding a K_i_ of c-di-GMP for CobB of 32.27 μM (**Fig. 2b**). Thus, c-di-GMP noncompetitively inhibits the deacetylation activity of CobB, suggesting that the binding region of c-di-GMP on CobB is not in the catalytic pocket.

To further confirm the inhibition of c-di-GMP on CobB activity, we sought to alter the endogenous levels of c-di-GMP and test the deacetylation activity of CobB *in vivo*. In *E.coli*, YdeH^14^ is the major DGC that produces c-di-GMP. Thus, we examined an *E.coli* strain with overexpressed YdeH (y*deH*^+^) and a strain expressing a dysfunctional mutant YdeH (*ydeH*^G206A,G207A^)^14^ to produce strains with high and low levels of c-di-GMP, respectively. To monitor the activity of CobB, we examined the acetylation level of endogenous Acs to which a chromosomal C-terminal FLAG-tag was attached. We also examined a series of CobB-depleted strains, namely Δ*cobB*, Δ*cobB::cobB* and Δ*cobB::cobB ydeH*^+^, to further investigate this interaction. As expected, high levels of c-di-GMP were clearly observed for strains with YdeH^+^ **(Fig.2c)**. In these strains, we found a significant reduction of deacetylation of Acs **(Fig.2d)**. We found that this deacetylation of Acs is abolished by the deletion of CobB (Δ*cobB*), and that this could be recovered by putting back CobB (Δ*cobB::cobB*). Thus, these results indicate that c-di-GMP increases the acetylation levels of Acs in a CobB-dependent manner.

It is known that CobB activates Acs through deacetylation of K609, and Acs is responsible for the synthesis of acetyl-CoA, which is essential for cell growth^22^. It is also known that both acetate and propionate can serve as a donor for acetyl-CoA for cell growth^33, 34^ **(Fig.2e)**. To further confirm the regulatory role of c-di-GMP on acetyl-CoA synthesis, all of the aforementioned strains **(Fig.2f)** were cultured using acetate or propionate as the sole carbon source. For the CobB deficient cells (Δ*cobB*), we observed an obvious growth defect, consistent with a lack of activated Acs. Similarly, overexpression of YdeH significantly inhibited cell growth in the *ydeH*^+^ and Δ*cobB::cobB ydeH*^+^ strains. These results are consistent with an inhibition of CobB by high levels of c-di-GMP **(Fig.2f, Supplementary Table 1)**. We also found that the *ydeH*^G206A,G207A^ strain exhibited a similar growth as that of the WT strain. The inhibition of growth in the strains with *ydeH*^+^ was similar at concentrations of acetate or propionate of 10 mM or 30 mM **(Supplementary Fig. 5, Supplementary Table 1)**. Thus, in addition to confirming the c-di-GMP inhibition of CobB *in vivo*, these results strongly suggest that c-di-GMP is a physiologically relevant effector of the regulation of acetyl-CoA biogenesis through the inhibition of the deacetylation activity of CobB.

### c-di-GMP globally affects CobB-dependent deacetylation in *vivo*

To determine whether c-di-GMP could affect CobB-dependent deacetylation in a global setting, we applied Stable Isotope Labeling with Amino acids in Cell culture (SILAC) coupled with MS to quantitatively compare the levels of protein acetylation in WT and *ydeH*^+^ cells. Previously, Weinert *et al*.^34^ identified 366 CobB regulated acetylation sites when *E.coli* was cultured in M9 media supplemented with 0.2% glucose. In addition, Cerezo *et al.^35^* identified 283 acetylation sites in *E.coli* under carbon-limited conditions. Since the acetylome of *E.coli* varies significantly under different culture conditions, to facilitate comparisons with previous work, we adopted the procedure developed by Weinert *et al*.^34^ with slight modifications (see Methods). A total of 802 acetylation sites (**Data set S1**) were identified, of which 107 (**Data set S2**) exhibited enhanced acetylation upon overexpression of YdeH **(Fig. 3a, b, supplementary Fig. 6)**. Of the CobB regulated acetylation sites identified by Weinert *et al*.^34^ (**Data set S3**), 43 were discovered in our study, 28 of which were among those that exhibited greater acetylation in *ydeH*^+^ **(Fig. 3c, data Set S4)**. Hence, CobB regulated sites are enriched in *ydeH*^+^ cells. Furthermore, we mapped the 107 c-di-GMP upregulated sites to 87 proteins and found that 42 of these proteins overlap with the 271 CobB regulated proteins identified previously^34^ **(Fig. 3d, data Set S5)**. We also compared the functional categories of the c-di-GMP upregulated proteins with the CobB regulated proteins and found that these two sets of proteins are highly similar in several classifications **(Supplementary Fig. 7)**. We note that slight differences in our procedure from that described by Weinert *et al*. (particularly the absence of fractionation before MS analysis) may explain the lower numbers of acetylation sites identified here compared to the earlier work (802 vs. 3,680). Nevertheless, our data strongly indicate that c-di-GMP regulated acetylation is closely related to CobB dependent deacetylation. Thus, these results suggest that c-di-GMP globally affects CobB-dependent protein deacetylation *in vivo*.

**Figure 3.**
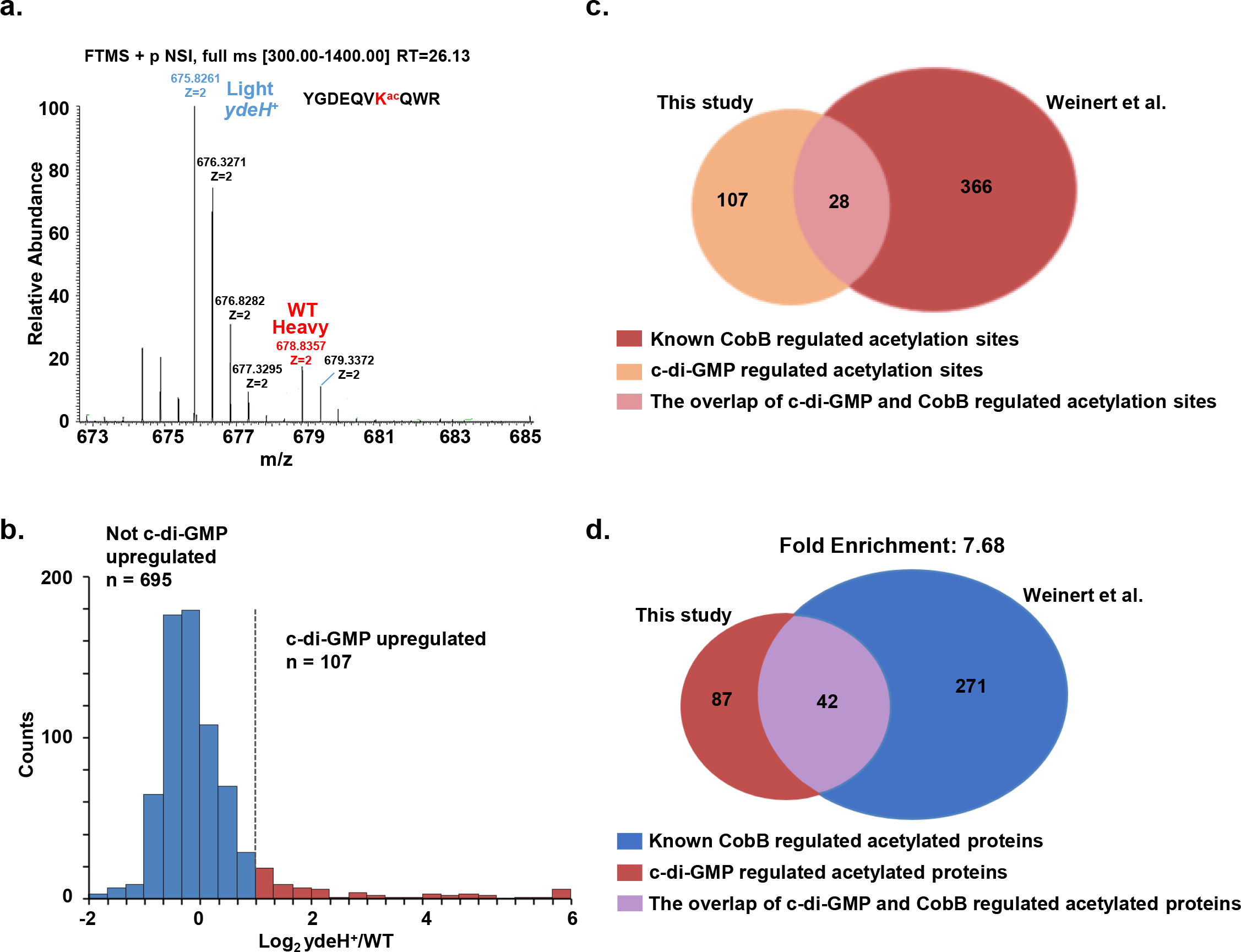
c-di-GMP globally affects CobB-dependent deacetylation. c-di-GMP affects *E.coli* acetylation. Two strains, *i.e*., WT and *ydeH*^+^ were included for quantitative acetylation analysis using SILAC-MS. **(a)** The mass spectra show light (*ydeH*^+^) and heavy (WT) signal for a representative peptide. **(b)** Histogram shows the SILAC ratio distribution of acetylation sites in *ydeH*^+^ cells compared to that of the WT cells. The levels of 107 acetylated peptides were upregulated in *ydeH*^+^. **(c)** The pie charts show the overlap of the c-di-GMP regulated acetylation sites and the known CobB regulated acetylation sites reported by Weinert *et al*. **(d)** The charts show the overlap of c-di-GMP regulated acetylation proteins and the known CobB regulated acetylation proteins reported by Weinert *et al*.

### Mutagenesis and binding studies of the CobB and c-di-GMP interaction

There is an additional 37 aa N-terminal tail in CobB compared to that of CobB_S_^21^ (**Fig. 4a**). Streptavidin blotting assays showed that truncation of the N-terminal fragment of CobB disrupts c-di-GMP binding, suggesting that residues 1-37 of CobB are essential for c-di-GMP interaction (**Fig. 4b**). It is known that c-di-GMP binds to many of its effectors by Arg, Leu, Asp and Glu residues^36^. To determine the exact binding sites on CobB, we mutated all Arg, Leu, Asp and Glu residues to Ala within the CobB N-terminal domain. We found that only CobB mutants with R8A, R17A and E21A exhibited a weakened interaction with c-di-GMP **(Fig. 4c)**. We next performed ITC titration with these mutants and found that the *K_d_*values are 1.8 mM, 1.6 mM and 0.32 mM for CobB^R8A^, CobB^R17A^ and CobB^E21A^, respectively **(Fig. 4d)**, which is 15 to 83-fold lower than that of the wild type CobB **(Fig. 1c and supplementary Fig. 1d-f)**. Despite a loss of c-di-GMP binding, CobB^R8A^, CobB^R17A^ and CobB^E21A^ displayed similar deacetylase activity as CobB. Importantly, addition of c-di-GMP did not inhibit the activity of these mutants **(Fig. 4e)**. Collectively, these data suggest that R8, R17 and E21 are important for c-di-GMP binding but do not directly participate in the catalytic activity of CobB.

**Figure 4.**
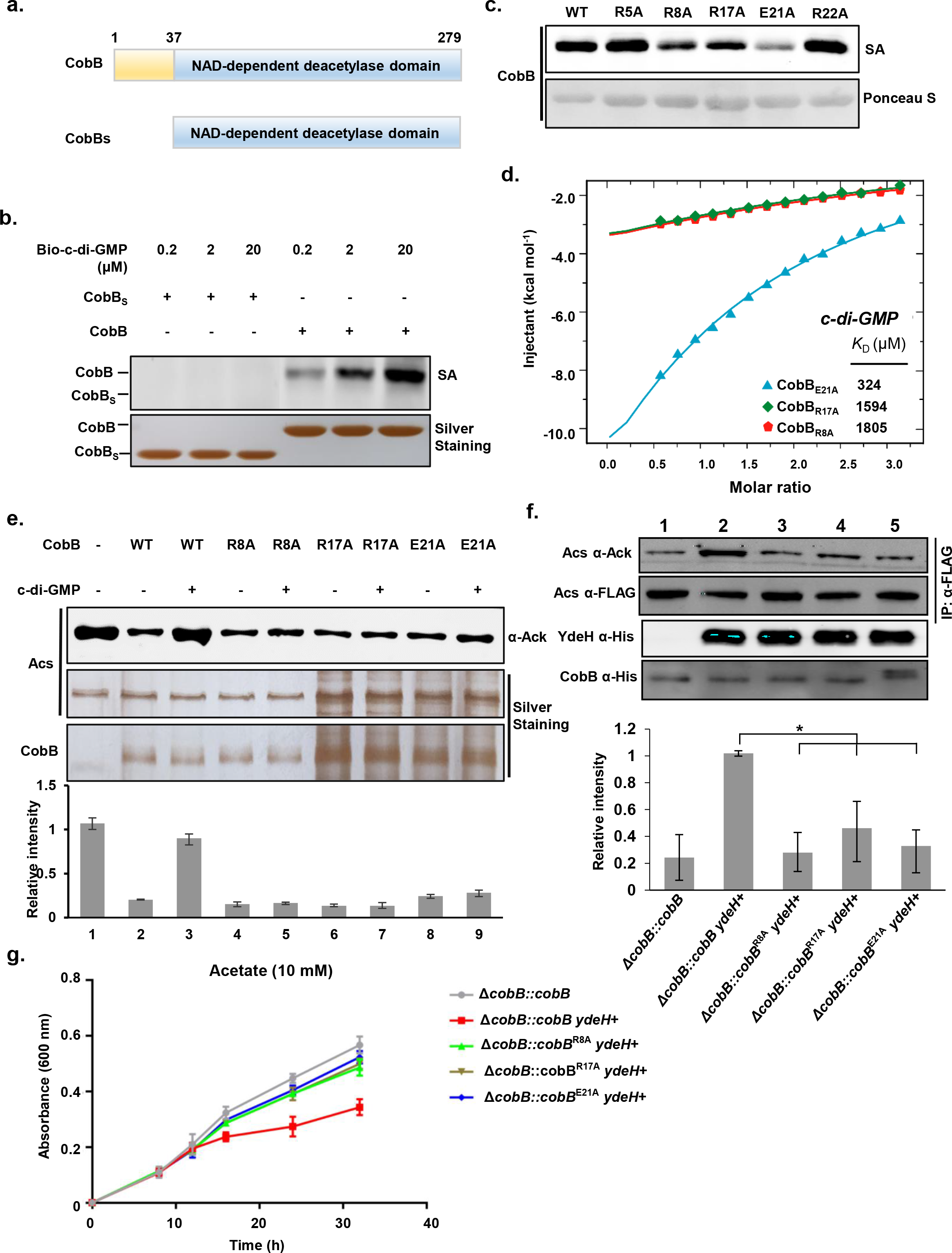
Determination of the binding sites of c-di-GMP on CobB. **(a)**CobB domain architecture shows the N-terminal domain (colored yellow). While for the truncation version of CobB, *i.e*., CobB_S_, there is no N-terminal domain. **(b)** CobB_S_ lost the binding with c-di-GMP. Both CobB and CobB_S_ were incubated with bio-c-di-GMP. After UV cross linking, the reactions were subjected for streptavidin blotting. **(c)** Streptavidin blotting assays for WT and CobB mutants. It is known that c-di-GMP tends to bind residues Arg (R) and Glu (E). We individually mutated all the R and E residues to pinpoint the binding sites. The c-di-GMP binding assays were carried out with all these mutated CobB, the bindings were visualized by streptavidin. The loadings were monitored by Ponceau S staining. **(d)** ITC assay to measure the binding kinetics of c-di-GMP and three CobB mutants. **(e)** *In vitro* deacetylation assay of the three CobB mutants using Acs as substrate with three replicates. **(f)** The deacetylation activity of CobB mutants were monitored using endogenous Acs as substrate *in vivo*. (three preparations; two-tailed Student’s t-test, \**P* < 0.05) **(g)** The growth curves of the CobB mutanted strains were determined. The growth was measured at 8, 12, 16, 20, 24 and 32 h with three replicates.

To further validate the specific binding of c-di-GMP to CobB, we constructed strains of these three CobB mutants, *i.e*., Δ*cobB::cobB*^R8A^*ydeH*^+^, Δ*cobB::cobB*^R17A^ *ydeH*^+^ and Δ*cobB::cobB*^E21A^ *ydeH*^+^ based on an *E.coli* strain that includes the chromosomal Flag-tagged Acs. These strains were cultured with acetate or propionate and the c-di-GMP concentrations were measured (**Supplementary Fig. 8a)**. To monitor the *in vivo* activity of CobB, we examined the acetylation of endogenous Acs. As expected, the acetylation levels of Acs were significantly lower in the strains with these CobB mutants, compared with that in the Δ*cobB::cobB ydeH*^+^ strain **(Fig. 4f)**. Additionally, these CobB mutant strains showed similar growth rates as Δ*cobB::cobB*, but higher growth rates than Δ*cobB::cobB ydeH*^+^ **(Fig. 4g, supplementary Fig. 8b-d)**. Hence, these results provide *in vivo* evidence for the role of these residues in CobB in the binding of c-di-GMP.

### The binding and inhibition of the deacetylation activity of CobB by c-di-GMP is conserved among prokaryotes

CobB is a Sir2 homolog that is highly conserved in prokaryotes. We thus hypothesized that the binding and inhibition of c-di-GMP observed with CobB from *E.coli* is the same for the CobB homologues in other prokaryotes. To test this hypothesis, CobB protein sequences from a series of highly diverse bacteria were aligned **(Supplementary Fig. 9)**. We found that the c-di-GMP binding region is reasonably well conserved in these bacteria **(Fig. 5a)**. Examining *S. typhimurium* CobB as an exemplary member of this conserved set, we found that CobB^*S. typhimurium*^ binds to c-di-GMP, but not to c-di-AMP and cGMP (**Fig. 5b**). Furthermore, we determined the affinity of *S. typhimurium* CobB and c-di-GMP to be 15.5 μM **(Supplementary Fig. 10)**, and that the CobB^*S.typhimurium*^ deacetylation of Acs could be clearly inhibited by c-di-GMP, but not cGMP and c-di-AMP **(Fig. 5c)**.

**Figure 5.**
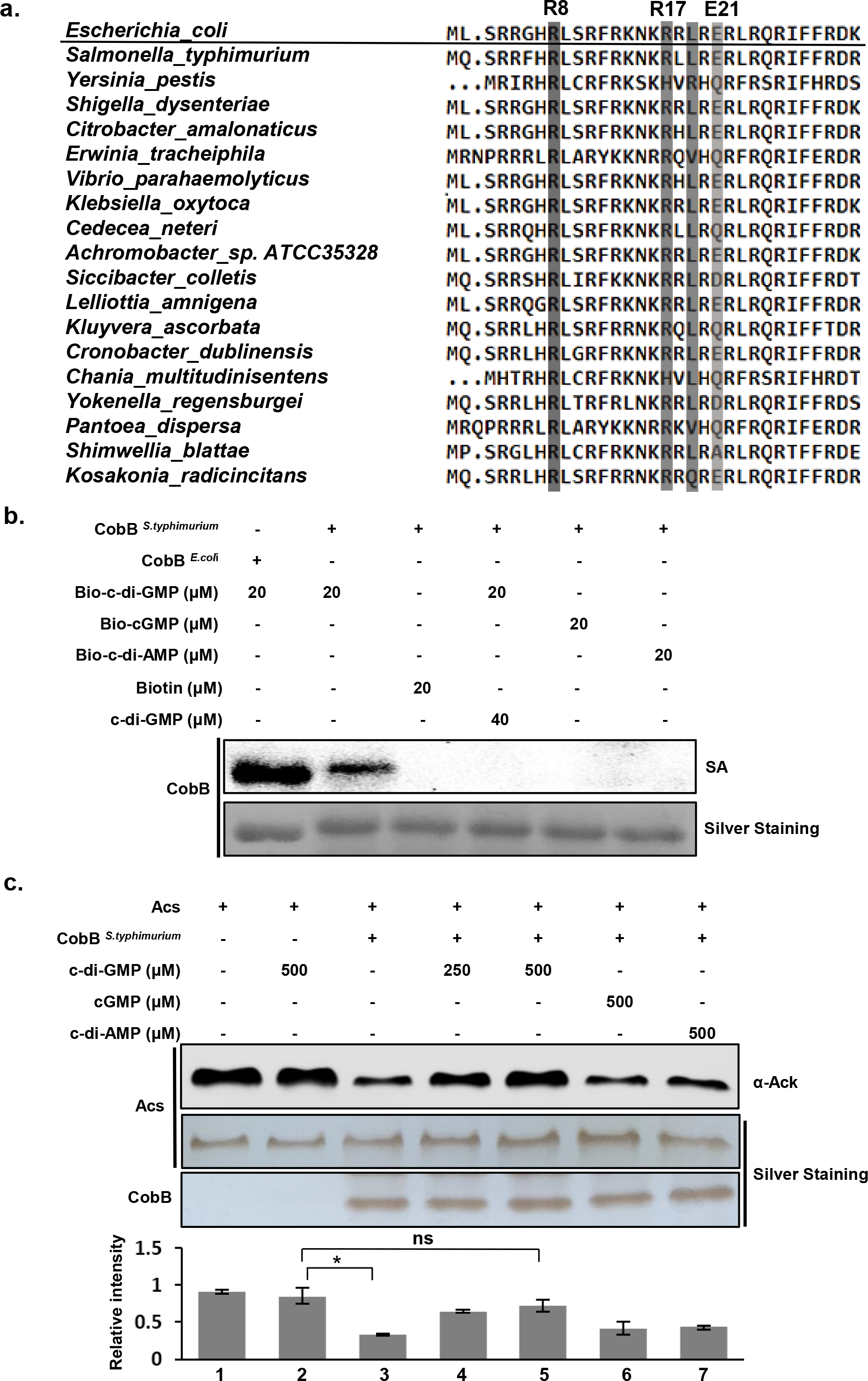
c-di-GMP binds CobB and inhibits its activity is conserved in prokaryotes. **(a)**CLUSTALW alignment of the binding motif of c-di-GMP and CobB. Residues involved in c-di-GMP binding R8, R17 and E21 were boxed (Gray) and the depth of color indicated the degree of conservative of the residues. **(b)** c-di-GMP binding assay of *E.coli* and *S. typhimurium* CobB. The *E.coli* and *S. typhimurium* CobB were incubated with bio-c-di-GMP, same amount of biotin, cGMP and c-di-AMP were included as controls. **(c)** c-di-GMP inhibits *S. typhimurium* CobB’s deacetylase activity *in vitro*. CobB^*S. typhimurium*^ activity assay was performed using *E.coli* Acs as the substrate. The acetylation level of Acs was detected by the pan anti-acetyl antibody. (three preparations; \**P* < 0.05, two-tailed Student’s t-test).

### CobB deacetylates YdeH on K4 to activate its DGC activity

Protein acetylation is one of the most abundant post-translational modifications in bacteria (and eukaryotes), with hundreds of acetylated proteins already identified in *E.coli*^37, 38^. We speculated that some of the proteins involved in c-di-GMP metabolism are endogenously acetylated and could be deacetylated by CobB. We selected 4 DGCs (YdeH, YaiC, YegE and YeaJ), 2 PEDs (YahA, DosP), and 3 GTPases (Era, YihK and YihI) from *E.coli* to test this possibility. We found that the 2 DGCs (YdeH, YegE) and 2 GTPases (Era, YihK) were acetylated **(Fig. 6a)**. Further, treating these proteins with CobB, we found that YdeH (DGC) and Era (GTPase) could be effectively deacetylated by CobB **(Fig. 6a)**. To determine the residue(s) of YdeH targeted for deacetylation by CobB, both acetylated YdeH and CobB-treated deacetylated YdeH were subjected to mass-spectrometry analysis. In this way, YdeH K4, K170, K277 were determined as the sites that could be deacetylated by CobB **(Fig. 6b, supplementary Fig. 11)**. To validate these sites, we mutated all 8 Lys residues of YdeH to Ala individually, and found that only the YdeH K4 mutant exhibited a significantly decreased acetylation level **(Fig. 6c)**. These results indicate that K4 is the dominant acetylated site on YdeH. In addition, two other acetylation sites were also identified on Era (K171) **(Supplementary Fig. 12a)** and YegE (K936) **(Supplementary Fig. 12b)**.

**Figure 6.**
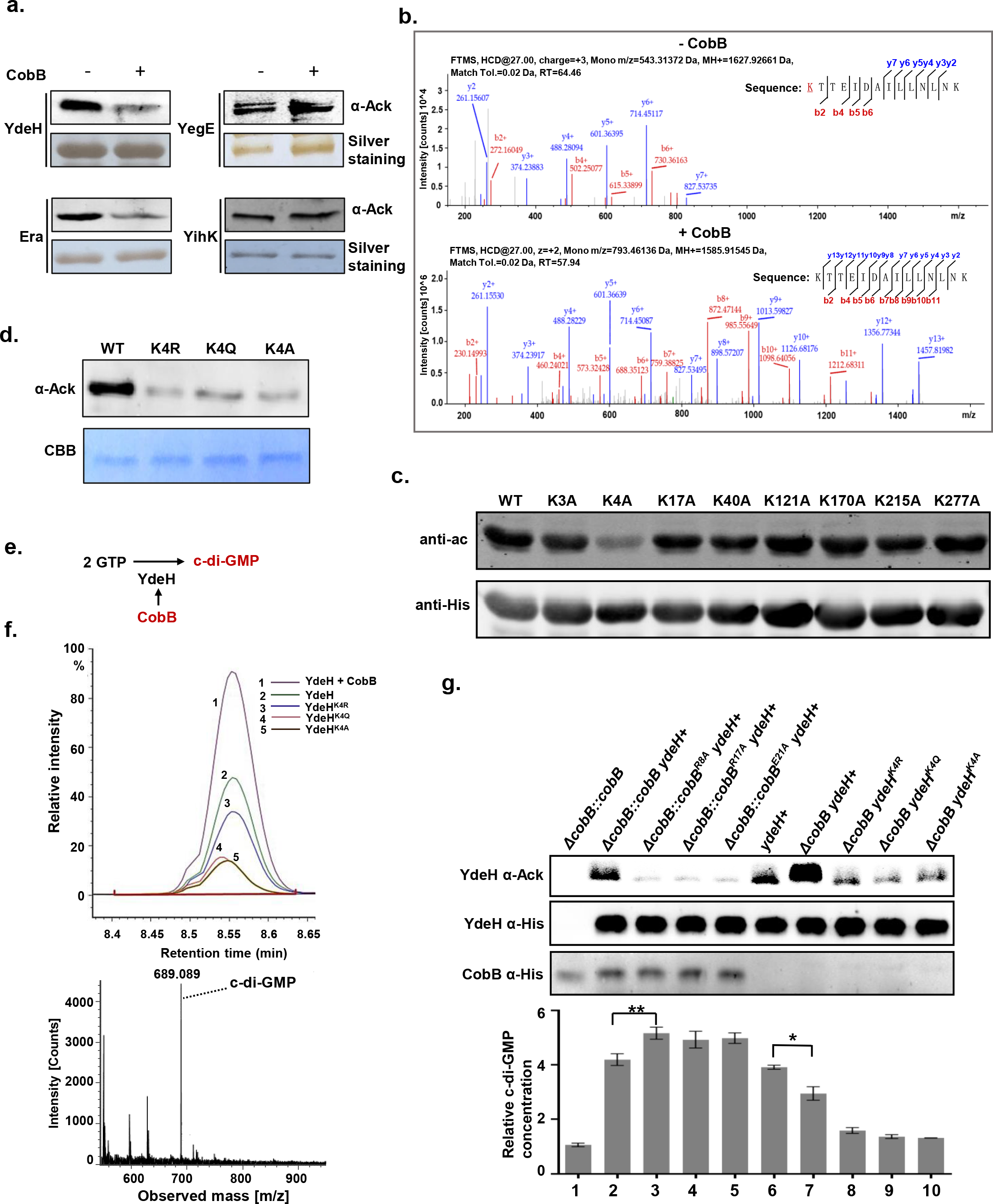
CobB up-regulates the level of c-di-GMP through deacetylation of the major diguanylate cyclase YdeH in *E.coli*. **(a)** The acetylation levels of four c-di-GMP related enzymes of *E. coli* with or without CobB treatment was monitored by the pan anti-acetyl antibody. **(b)** Mass spectrometry (MS) analysis to determine the acetylation site/s of YdeH that could be deacetylated by CobB. The affinity purified YdeH with or without CobB treatment were subjected for MS analysis. K4 was discovered as the deacetylation site. **(c)** Mutagenesis of YdeH confirmed that K4 is the major acetylation site. All the 8 Lys residues of YdeH were mutated to Ala individually. The acetylation was visualized using the anti-acetyl antibody. **(d)** Mutagenesis of YdeH K4 confirms the acetylation of K4. Three single YdeH mutants, K4R, K4Q and K4A were constructed. **(e)** CobB modulates the synthesis of c-di-GMP through deacetylating YdeH. **(f)** The DGC activity of the three mutants along with WT YdeH and CobB treated WT YdeH were analyzed using UPLC-IM-MS *in vitro* with three replicates, the peak areas represent the amounts of c-di-GMP in these samples. **(g)** YdeH’s acetylation levels and c-di-GMP concentrations were measured with CobB mutants and YdeH mutants. The acetylation level of purified YdeH was detected by the pan anti-acetyl antibody and the protein levels of YdeH and CobB were determined by the anti-His antibody. (three preparations; \**P* <0.05, \*\**P* < 0.01, two-tailed Student’s t-test).

To verify the functional role of YdeH K4 acetylation, we mutated the Lys to Arg (YdeH^K4R^), Gln (YdeH^K4Q^) or Ala (YdeH^K4A^). Western blotting showed that these mutations significantly reduced the level of acetylation of YdeH compared to that of WT YdeH **(Fig. 6d)**. Since YdeH is the major DGC in *E.coli*, we then set to test whether CobB deacetylation could affect the DGC activity of YdeH using UPLC-IM MS (**Fig. 6e**). Upon CobB treatment, WT YdeH showed a 2-fold increased production of c-di-GMP, while the activities of YdeH^K4R^, YdeH^K4Q^ and YdeH^K4A^ were 69.5%, 29.5%, and 27.1% of that of the WT, respectively **(Fig. 6f)**. Unexpectedly, the mutant YdeH^K4R^ that structurally resembles the deacetylation form of YdeH also exhibited a lower DGC activity. Evidently, the “K” to “R” mutation at residue 4 fails to fully mimic the deacetylation status of this site. Nonetheless, these results suggest that K4 is critical for the DGC activity of YdeH.

To test whether CobB can regulate the c-di-GMP levels *in vivo*, we constructed several *ydeH*^+^ and Δ*cobB ydeH*^+^ strains, including those with CobB mutants or with YdeH mutants, and then determined the YdeH acetylation and c-di-GMP levels. We found that the acetylation level of YdeH in *ydeH*^+^ is lower than that of Δ*cobB ydeH*^+^ and the c-di-GMP level of *ydeH*^+^ is significantly higher than that of Δ*cobB ydeH*^+^ **(Fig. 6g)**. These results confirm that CobB can regulate the DGC activity of YdeH through deacetylation. The strains with the YdeH mutants exhibited the lowest DGC activity, confirming that K4 is critical to the DGC activity of YdeH **(Fig. 6g).** Interestingly, the strains with the CobB mutants exhibit lower acetylation levels than WT CobB **(Fig. 6g)**. These results indicate that c-di-GMP does not affect the activity of the CobB R8A, R17A and E21A mutants in *vivo*. Still, the CobB mutants exhibited higher c-di-GMP levels, further confirming that CobB promotes the DGC activity of YdeH through deacetylation.

### CobB enhances the solubility/stability of YdeH through deacetylation

We found that the acetylation level of endogenous YdeH is significantly decreased in WT cells compared to that of CobB-deficient cells **(Fig. 7a)**. In order to better characterize the endogenous stoichiometry of YdeH K4 acetylation, we used the absolute quantification (AQUA) method^39, 40^ to quantify the level of the K4 acetylated peptide. As shown in **Fig. 7b**, the YdeH K4 acetylation stoichiometry was 1.3 ± 0.2 % for the CobB deficient cells, while it is undetectable for the WT cells.

**Figure 7.**
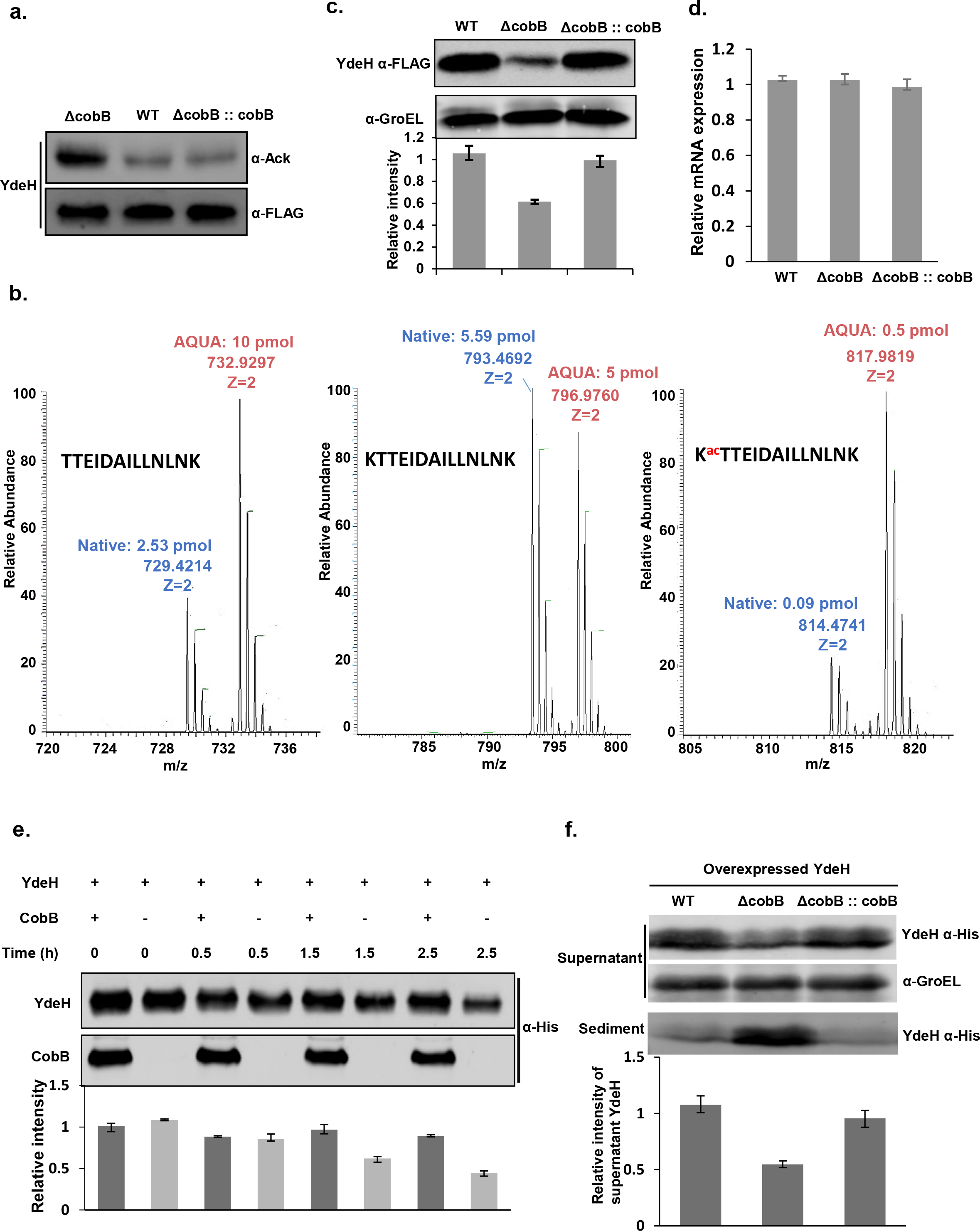
YdeH is endogenous acetylated and this acetylation is regulated by CobB. **(a)**The endogenous YdeH acetylation level of 3 *E.coli* strains. A 3xFLAG tag was chromosomally inserted at the 3’-end of Acs coding sequence. 3xFLAG-tagged YdeH was enriched by an anti-FLAG antibody and the acetylation levels were determined by the pan-Ack antibody. **(b)** The mass spectra show AQUA quantification of the endogenous acetylation of YdeH K4 using AQUA peptides. **(c)** The level of the endogenous YdeH. The protein levels were determined by the anti-FLAG antibody and GroEL was applied as the loading control. The bar graph showed the quantitation of the protein levels of YdeH with three replicates. **(d)** The relative mRNA repression of endogenous YdeH. **(e)** The stability of YdeH with or without CobB treatment. These samples were separated at 0.5 h, 1.5h and 2.5 h and the protein levels were determined by the anti-His antibody. The bar graph showed the quantitation of the protein level of YdeH with three replicates. **(f)** The level of the overexpressed YdeH in the supernatant and sediment. The protein levels were determined by the anti-His antibody and GroEL was applied as the loading control. The bar graph showed the quantitation of the protein level of supernatant YdeH with three replicates.

Since it is known that acetylation can lead to protein degradation *in vivo*^41^, we speculated that acetylation of YdeH may affect its stability. Indeed, we found that the level of endogenous soluble YdeH deceased 40% in CobB deficient cells as compared to that of the WT cells (and also the CobB recovered cells) in exponential growth **(Fig. 7c)**, and more significantly, a decrease of 70% soluble YdeH was observed for stationary growth **(Supplementary Fig. 16c)**. Furthermore, for stationary growth, a significant amount of precipitated YdeH was observed for CobB deficient cells, but not for WT and CobB recovered cells **(Supplementary Fig. 16c)**. We then examined the protein stability of YdeH mutants *in vitro* and found that YdeH^WT^ and YdeH^K4Q^ are less stable than YdeH^K4A^ and YdeH^K4R^ **(Supplementary figure 13)**. In addition, we also noticed that the level of soluble YdeH following the addition of CobB is 2-fold of that in the absence of CobB **(Fig. 7e)**.

Additionally, our data show that the c-di-GMP increase by 36% in WT cells than CobB deficient cells *in vivo* under exponential growth when YdeH was overexpressed **(Fig. 6g)**. To understand the underlying mechanism, we then determined the level of YdeH when it was overexpressed in WT cells and CobB deficient cells. The results showed that the soluble YdeH level decreased ~45% in CobB deficient cells as compared to that of the WT cells **(Fig. 7f)**, which is similar to that of the endogenous YdeH under exponential growth (**Fig. 7c**). We also measured the YdeH level in the sediment and observed significant amount of precipitated YdeH in CobB deficient cells as compared to that of WT and CobB recovered cells **(Fig. 7f)**.

Thus, it appears as though YdeH K4 acetylation lead to protein aggregation/precipitation by reducing its solubility/stability **(Fig. 8a)**. In WT cells, the acetylation of YdeH is removed by CobB and retained in soluble state, with little or undetectable protein sediment **(Supplementary Fig. 16c)**. In CobB deficient cells, more acetylated YdeH will precipitate with both acetylated and/or deacetylated YdeH because of the lack of CobB deacetylation. Additionally, this model also explains why YdeH K4 may be frequently acetylated but cannot accumulate to high level *in vivo*. It is known that acetylation on specific sites promote protein aggregation^42, 43^. For example, K280 acetylation causes the aggregation of tau. And tau promotes neuronal survival, thus the acetylation induced tau aggregation is pathologically significant. Interestingly, tau K280 belongs to a double lysine motif, *i.e*., ^275^VQIINKK^281^, and YdeH K4 belongs to a similar double lysine motif, *i.e*., ^1^MIKK^4^. Thus, it is possible that the underlying mechanism of lysine acetylation induced protein aggregation of YdeH may be similar to that of tau. However, because of the possible complexity, the detailed mechanism is yet to be discovered.

**Figure 8.**
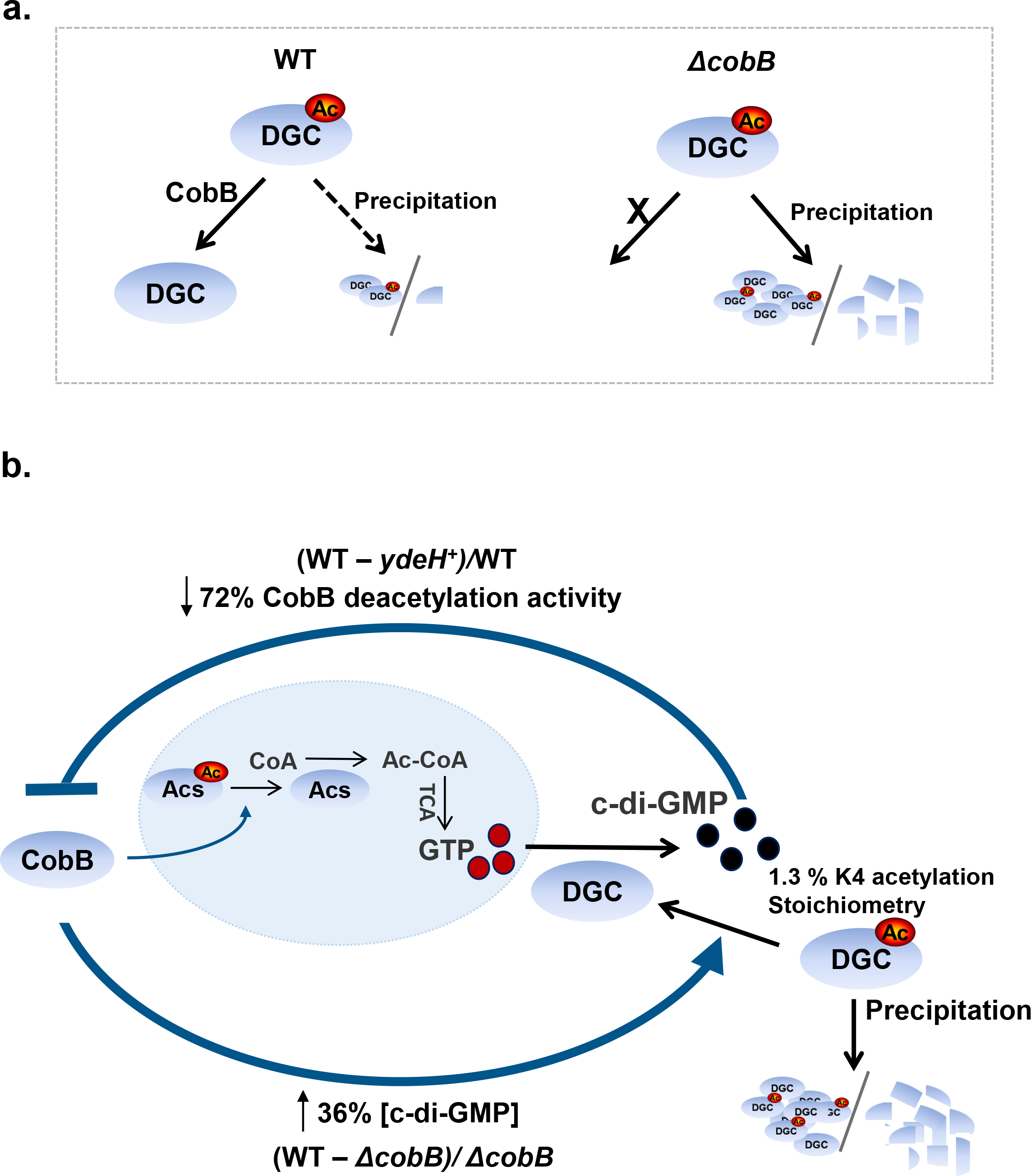
The regulating model of c-di-GMP and CobB interplay. **(a)**YdeH K4 acetylation reduce YdeH’s stability and cause precipitation. **(b)**The overview of c-di-GMP and CobB interplay. c-di-GMP inhibits the deacetylation activity of CobB and then regulates Ac-CoA synthesis. In another direction, CobB activates YdeH’s DGC activity through deacetylation to prevent the precipitation of YdeH, and also activate YdeH. In addition, Ac-CoA modulates GTP, the precursor of c-di-GMP generation through TCA. Quantitatively, c-di-GMP abolishs 72% of CobB’s decetylation activity through direct binding of CobB, and CobB promotes c-di-GMP level to 36% more through deacetylation of YdeH.

## DISCUSSION

c-di-GMP is a key secondary messenger in prokaryotes. CobB is the first, and major, protein deacetylase identified in prokaryotes. Herein, we discovered that c-di-GMP specifically binds to CobB and inhibits its deacetylase activity, and down-regulates the cellular concentration of acetyl-CoA through modulation of the acetylation levels of Acs. Interestingly, we also found the major *E.coli* DGC, YdeH, is endogenously acetylated, and CobB enhances the solubility/stability of YdeH, and activates the DGC activity of YdeH, through deacetylation. Thus, we established a negative feedback regulatory loop between c-di-GMP biogenesis and CobB dependent protein deacetylation.

### The binding region and kinetics of CobB and c-di-GMP

The most well-known motif of c-di-GMP binding is the EXLXR motif^44^. Interestingly, the c-di-GMP binding site on CobB contains RXLXE, which is precisely EXLXR in reverse. As expected in the EXLXR motif, both R17 and E21 are critical for c-di-GMP binding, while the L19 residue does not directly bind to c-di-GMP in the RXLXE motif. This motif is located in the N-terminal tail of this protein that we found is responsible for its dimerization **(Supplementary Figure 3)**. Since many of the c-di-GMP binding proteins are dimers or tetramers^4, 31, 45^, a plausible explanation for this effect of c-di-GMP described here is that c-di-GMP interferes with the dimerization of CobB.

A concern with our model though is that the K_d_ and K_i_ of c-di-GMP for CobB measured *in vitro* are larger than the sub- to micro-molar concentrations of c-di-GMP generally observed *in vivo*^46^. While this may be owing to presently unidentified molecular factors that reduce this affinity *in vivo*, we believe that this discrepancy could be explained by the spatially and temporally uneven distribution of c-di-GMP in the bacteria. Indeed, direct measurement of the concentration of c-di-GMP in bacteria using a FRET biosensor has revealed a wide range of concentrations within individual cells^47^, including some resolution-limited locations (~200 x 200 x 1000 nm^3^) that exhibit a local concentration significantly greater than 1 μM. We note that this concentration corresponds to only 25 molecules within this volume. Further, it is well known that the bacterial cytoplasm is extremely crowded^48^, and recent work has shown that, as a result, effectively, there are caging effects on the free diffusion of particles^49^. A single molecule of c-di-GMP within a “cage” of only 20 x 20 x 100 nm^3^ is at a concentration of 40 μM. Thus, we speculate that it is indeed physically possible that the local concentration of c-di-GMP could exceed the measured affinity constants for CobB in the WT bacteria. In fact, owing to the mechanism described here, the negative regulatory loop involving CobB and YdeH may be principally responsible for keeping the local (and thus global) cytosolic concentration to the observed maximal levels.

### c-di-GMP regulates physiological functions through inhibiting of CobB deacetylation

As the most important substrate of CobB, Acs is responsible for the synthesis of acetyl-CoA, which controls cell energy metabolism and global protein acetylation, and affects cell growth and proliferation^50-52^. Our results indicate that c-di-GMP can lower the concentration of acetyl-CoA *in vivo* through inhibiting CobB and increasing the acetylation level of Acs. Previous studies have identified several possible overlapping effects of c-di-GMP and acetyl-CoA. For example, c-di-GMP affects the expression of acetate kinase (AckA) through the binding of the transcription factor that regulates AckA transcription in *B. burgdorferi*^53^. In addition, acyl-CoA dehydrogenase, a key enzyme in acetyl-CoA metabolism pathways, is a genuine c-di-GMP effector in *B. bacteriovorus*^54^. Here, we strengthened the link between c-di-GMP and acetyl-CoA by showing, for the first time, that c-di-GMP directly modulates the biogenesis of acetyl-CoA.

c-di-GMP has emerged as a key regulator in the decision between motile and sedentary forms of bacteria^2^. Elevated c-di-GMP levels inhibit bacterial motility via effects on the flagella-associated protein^6^. Most recently, Nesper *et al*. found that c-di-GMP directly binds to the CheY-like regulators and tunes the bacterial flagellar motor^9^. According to our data, c-di-GMP can regulate the acetylation level of CheY via inhibition of the deacetylase activity of CobB. Thus, it is highly possible that c-di-GMP regulates bacterial motility through CobB-mediated regulation of CheY activity.

CobB is a member of the sirtuin protein family, which are highly conserved across prokaryotes to eukaryotes. Thus, it is possible that c-di-GMP may also bind human sirtuins and play critical functional roles in a variety of biological processes, such as host-pathogen interactions.

### CobB regulates the YdeH DGC activity *in vitro*

The overexpressed YdeH was used for DGC activity assay *in vitro*. We determined the K4 acetylation stoichiometry of the overexpressed YdeH following the same AQUA procedure that we used for the endogenous YdeH. The results showed that the overexpressed YdeH K4 acetylation stoichiometry is 28.7 ± 4.9% **(Supplementary Fig. 14)**. Assuming that all the K4 acetylation could be deacetylated by CobB, we could thus expect a ~1.4-fold activity increase upon CobB deacetylation. But it is still could not fully explain the 2-fold activity increase upon CobB deacetylation **(Fig. 6f)**. To this end, we sought to determine, with a simple model, whether the loss of stable YdeH could completely account for the 2-fold difference in c-di-GMP. In particular, based on the stability data (**Fig. 7e**), we considered the case in which the difference in c-di-GMP is completely owing to the decrease in soluble protein. Assuming that the substrate is not limiting throughout the assay and that steady state is achieved after a very short time, the lower amount of c-di-GMP produced would simply be a result of the lower amount of soluble protein at any time point. We found that the decrease in soluble protein is well-described by a single exponential **(Supplementary Fig. 15)**

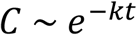

where *C* is the concentration of YdeH and *k* is the rate of the protein loss, determined to be 0.49 hr^-1^ in the fit. Thus, the fold-increase, *F*, expected for the fully stable (CobB treated) YdeH relative to the more unstable version can be calculated (for the 2 hrs assay) by

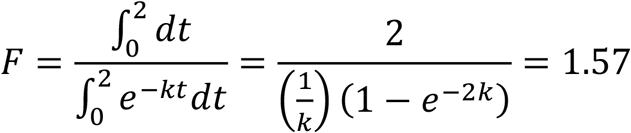

with the aforementioned value of *k*. Thus, simple loss of the active protein would result in a 1.57-fold difference in c-di-GMP. Since we observed a 2-fold difference, we conclude that the loss of soluble protein can account for ~78% (1.57/2) of the difference measured with the acetylated and deacetylated YdeH. Hence, a difference in the inherent activity between acetylated-YdeH and deacetylated-YdeH is needed to fully account for observed difference in c-di-GMP produced.

We noticed that the K4 acetylation stoichiometry of overexpressed YdeH is much high that of endogenous YdeH. The endogenous acetylation stoichiometry of YdeH K4 was measured using the exponential growth of *E. coli*. While as a common practice, when overexpress a protein in *E. coli*, the strain is usually induced at exponential growth, and then cultured for another ~2 hours for fast protein producing. At the time of protein purification, the strain is already at stationary phase. Thus, it is possible that the big difference of acetylation stoichiometry between the overexpressed YdeH and the endogenous one is at least partially due to the difference of the growth phases.

To test this possibility, we measured the YdeH K4 acetylation stoichiometry using *E. coli* of stationary growth. The results showed that the YdeH K4 acetylation stoichiometry is 1.1 ± 0.3% in WT cells and 4.2 ±0.5% in CobB deficient cells **(Supplementary Fig. 16a-b)**. For CobB deficient cells, the YdeH K4 acetylation stoichiometry is 3-fold(4.2%/1.3%) in stationary growth than that of exponential growth. At least, this data could partially explain the stoichiometry gap between the endogenous and overexpressed YdeH.

### CobB regulates the YdeH DGC activity *in vivo*

It is known that the expression of YdeH is tightly regulated by the RNA binding protein CsrA and the the *csrA* mutant can increase the transcription of YdeH in 15-fold and another DGC YcdT in 45-fold^55^. Thus, to artificially generate a condition of YdeH high expression for functional analysis, the *csrA* mutant has often been applied^56^. However, the *csrA* mutation can also activate the expression of another DGC, *i.e*., YcdT, which could cause high c-di-GMP background when studying YdeH. So, we chose to overexpress YdeH in *E. coli* to study the regulation role of CobB to YdeH inside the cell. Our data show that the c-di-GMP increase by 36% in WT cells than CobB deficient cells *in vivo* under exponential growth when YdeH was overexpressed **(Fig. 6g)**, which is consistent with the observed 45% decrease of soluble YdeH level in CobB deficient cells than that of WT cells **(Fig. 7f)**. Hence, the 36% lower c-di-GMP in CobB deficient cells is mainly due to the instability/precipitation of YdeH.

Though it is clear now that the major functional consequence of K4 acetylation is the decreasing of YdeH stability, the direct activity change upon acetylation and deacetylation by CobB may still play important role. Our results showed that YdeH^K4A^, which has comparable stability with WT YdeH, also exhibited a loss ~50 % of its DGC activity **(Fig. 6f)**. It is possible that the YdeH K4 acetylation may significant affect the structure of YdeH. To understand the mechanism, we proposed a working model that illustrates how CobB could regulate the structure of YdeH, based on an analysis of the atomic structure of YdeH^56^. It has been reported that dimerization and proper conformational rearrangement of the active center are required for optimal YdeH DGC activity **(Supplementary Fig. 17a)**. Reside K4 is located within a “hinge” region of YdeH and may potentially regulate the conformational rearrangement of the active center upon dimer formation **(Supplementary Fig. 17b-d)**. In the acetylated form of K4, the two GGDEF catalytic domains of YdeH are misaligned that may cause the aggregation and then lead to precipitate. **(Supplementary Fig. 17b)** Besides, this misaligned region could prevent productive ligation of the two GTP molecules, each captured by a GGDEF domain. According to our enzymatic studies, it is very likely that deacetylation of YdeH K4ac triggers the proper realignment of the GGDEF dimer so that the active center is rearranged to allow the formation of intermolecular phosphoester bonds between the two GTP substrates **(Supplementary Fig. 17c)**. Interestingly, the proposed model is close to the Zn-regulation mechanism in YdeH, *i.e*., inhibits YdeH’s DGC activity by hindering its dimerization^56^. Thus, at least in part, CobB may regulate the active/inactive switch of DGC activity of YdeH through deacetylation of YdeH on K4 **(Supplementary Fig. 17d).**

### The interplay between CobB and c-di-GMP

Our data show that c-di-GMP can significantly reduce of the decetylation activity of CobB through direct binding, and that the deacetylation of overexpressed YdeH by CobB leads to an increase of c-di-GMP by 36% *in vivo*. In addition, c-di-GMP is synthesized from GTP by DGC. CobB regulates the levels of acetyl-CoA through deacetylating Acs and then modulates the generation of GTP in the tricarboxylic acid (TCA) cycle. Thus, to a certain extent, the biogenesis of c-di-GMP could be activated by CobB through its effects on the TCA cycle. We also found that a GTPase Era and the DGC YegE are also endogenously acetylated. These results further demonstrate that the biogenesis of c-di-GMP is regulated by (de)acetylation.

The interplay between c-di-GMP and CobB may play important roles in bacteria. It is known that c-di-GMP participates in motility, biofilm formation and virulence, all of which are regulated by energy metabolism^57, 58^. We found that c-di-GMP modulates the bacterial energy metabolism through inhibiting the deacetylation activity of CobB, thus a connection between important bacterial processes and energy metabolism could be established. In addition, CobB is prominently implicated in the regulation of metabolism^59^ and c-di-GMP is involved in GTP metabolism and purine metabolism, suggesting a possible two-way regulation between c-di-GMP and CobB at the metabolic level.

Taken together, we established a novel feedback regulation loop between c-di-GMP and the deacetylation activity of CobB **(Fig. 8b)**. The findings of both directions (c-di-GMP inhibits CobB and CobB promotes c-di-GMP biogenesis) are novel. From this loop, we can envision a tightly regulated balance between the levels of c-di-GMP and the protein deacetylase activity of CobB. We strongly believe that our findings will facilitate future functional studies of both c-di-GMP and CobB-based regulation of protein acetylation.

## Methods

### Bacterial Strains, Plasmids, and CobB Mutant Construction

In this study, we used *E.coli* BW25113 as the wild type strain and the plasmid pSUMO10 for the overexpression of CobB, pET32a for the rescue experiment, and pCA24N for the overexpression of YdeH, Acs, CheY, NhoA, Era, YeaJ, YaiC, YahA, DosP, YihI, YegE and YihK. We performed the CobB mutations using the QuikChange^®^ Site-Directed Mutagenesis Kit (Agilent Technologies, Santa Clara, USA).

### *E.coli* Proteome Microarray Screening and Data Processing

The *E.coli* proteome microarrays were prepared as described previously^29^. The proteome microarrays were first blocked with blocking buffer (3% BSA in 0.1% Tween 20; TBST) for 1 h at room temperature. Bio-c-di-GMP was then diluted to 1 μM incubated on the microarray at room temperature for 1 h, and the same concentration of biotin was included as a negative control. The microarrays were next washed with TBST three times for 5 min each and then were incubated with Cy3-Streptavidin at a 1:1000 dilution (Sigma-Aldrich, Darmstadt, Germany) for 1 h at room temperature, followed by three washes with TBST for 5 min each. The microarrays were spun dry at 250 × g for 3 min and were scanned with a GenePix pro 6.1 microarray scanner to visualize and quantify the results. The raw data have been published in the Protein Microarray Database (www.proteinarray.cn) with the accession number, PMDE271. The auto-analysis tool in Protein Microarray Database (http://www.proteinarray.cn/index.php/analysis-toolkits) was used to process the microarray data.

### Preparation of c-di-GMP

It is known that c-di-GMP (BIOLOG Life Science Institute, Bremen, German) may form dimer or polymer, affects the binding with its effectors. Thus, we first determined the c-di-GMP monomer population. The OD_276_ and OD_289_ of c-di-GMP were measured using the cuvette mode of Nanodrop 2000c (Thermo Fisher Scientific, MA, USA) with a volume of 100 μL. To calculate the ratio of monomer of c-di-GMP (*P_mono_*)at various concentrations, we applied the equation (*p*_mono_ = 1.15 (*A*_276_/*A*_289_) − 1.64) developed by Gentner *et al.^60^* To eliminate the possible effect of c-di-GMP polymer, we heated the c-di-GMP solution in a water bath at 60°C for 1 h to depolymerize ~80% c-di-GMP to monomer before testing.

### Streptavidin blotting assay

In this assay, CobB (0.5 mg/mL, 16.13 μM) and 10 μM bio-c-di-GMP were incubated in the deacetylation buffer (50 mM Tris-HCl, 4 mM MgCl_2_, 50 mM NaCl, 50 mM KCl, 1 mM NAD^+^, pH 8.0) at 37°C for 1 h and the same amount of biotin, cGMP, c-di-AMP were included as negative controls. The samples were then UV-cross linked on ice with 10 min for 1.2 mJ/cm^2^ energy to further link the c-di-GMP to CobB. These samples were then analyzed by Western blotting. After incubation with IRDye 800-CW Conjugated Streptavidin (LI-COR Biosciences, Nebraska, USA) for 2 h, the membranes were washed with TBST three times and visualized with an Odyssey Infrared Imaging System (LI-COR Biosciences, Nebraska, USA). And the protein level was determined by Ponceau S staining is the (Sangon Biotech, Shanghai, China).

### ITC assay

In the ITC assay, c-di-GMP and the wild-type CobB and mutant CobB proteins were prepared in the titration buffer (20 mM Tris, 250 mM NaCl, 2 mM DTT, and pH 7.5). Protein concentrations were measured based on the UV 280nm absorption. The ITC titrations were performed using a MicroCal iTC200 system (GE Healthcare, Pittsburgh, USA) at 25°C. Each titration consisted of 17 successive injections (the first at 0.4 μL and the remaining 16 at 2.4 μL). The stock c-di-GMP, cGMP and c-di-AMP at 1.5 mM were titrated into wild-type or mutant CobB (0.1 mM) in the sample cells of 200 μL volume individually. c-di-GMP, cGMP and c-di-AMP of 1.5 mM were titrated into 200 μL titration buffer as controls for data processing. The resultant titration curves were processed using the Origin 7.0 software program (OriginLab) according to the “one set of sites” fitting model.

### CobB deacetylase activity assay

CobB (50 μg/mL, 1.6 μM) and c-di-GMP (at either 0.25 or 0.5 mM) were incubated in 20 μL deacetylation buffer (50 mM Tris-HCl, 4 mM MgCl_2_,50 mM NaCl, 50 mM KCl, 1 mM NAD^+^, pH 8.0) at 37°C for 0.5 h. For negative controls, we used 0.5 mM cGMP and c-di-AMP. The CobB substrates (4.16 μM Acs, 1.41 μM CheY, 0.62 μM NhoA) were then added and incubated at 37°C for 1.5 h. These proteins were analyzed by both silver staining and Western blotting. Membranes were incubated with a pan anti-acetyl antibody (Cell Signaling Technology, MA, USA, with a 1:1000 dilution) for at 4°C for 12 h and then incubated with an IRDye 800 secondary antibody at room temperature for 1 h. The membranes were visualized with an Odyssey Infrared Imaging System.

### Measuring the catalytic kinetics of CobB

CobB (9.3 μg/mL, 0.3 μM) was incubated with 0, 10, 20 and 80 μM c-di-GMP at 37°C for 20 min in 80 μL deacetylation buffer, and then 20 μL gradient concentrations acylated peptide (LEQIAELAGVSK^ac^TNLLYYFPSK)^32^ (1, 2, 4, 8, 16, 32, 50, 75, 100, 150 and 200 μM) were added and co-incubated at 37°C for 30 min. 100 mM HCl and 160 mM acetic acid were added to stop the reactions and spun for 10 min at 18,000 x g to separate the enzyme from the reactions. These samples were analyzed by HPLC. Briefly, samples were injected onto a C-18 column (AlltimaTM C18 4.6 x 250 mm) and analyzed by reversed-phase HPLC (Shimadzu, Japan). Solution A (0.065% trifluoroacetic acid in 100% water (v/v)) and solution B (0.05% trifluoroacetic acid in 100% acetonitrile (v/v)) were used in a gradient program (0.01 min with 5% Solution B, 25 min with 65% Solution B, 25.01 min with 95% Solution B, 31 min with 95% Solution B, 31.01 min with 5% Solution B, 40 min with 5% Solution B and stop in 40.01 min) with a flow rate of 1 mL/min. Peptides were detected at 220 nm wavelength. This assay was performed three preparations and V_max_, K_i_values were calculated by curve-fitting the plot using GraphPad Prism 6.

### Construction of an *E.coli* strain (BW25113) harboring chromosomal 3xFLAG-tagged Acs and 3xFLAG-tagged YdeH

*E.coli* strain (BW25113) harboring chromosomal 3xFLAG-tagged Acs and 3xFLAG-tagged YdeH were constructed using the Red recombination system^61^, as described previously^32^.

### Isolation and Quantification of c-di-GMP in *E.coli*

The c-di-GMP isolation methods were described previously^62^. Briefly, the total amount of *E.coli* cells with a 50 OD were harvested and re-suspended in 2 mL ddH_2_O. To extract intracellular c-di-GMP, 8 mL extract mixture of 50% methanol and 50% acetonitrile were added. We also added 1 μM cGMP as the reference. The mixture was incubated in boiling water for 10 min and then centrifuged at 10,000 rpm for 10 min. The supernatant was transferred to a new tube for freeze-drying. The freeze-dried pellet was re-suspended in 100 μL water containing 50% methanol. Samples were analyzed by UPLC-IM-MS (Ultra high-performance liquid chromatography coupled ion mobility mass spectrometry). UPLC-IM-MS was performed using a Waters UPLC I-class system equipped with a binary solvent delivery manager and a sample manager, coupled with a Waters VION IMS Q-TOF Mass Spectrometer equipped with an electrospray interface (Waters Corporation, Milford, USA) at the Instrumental Analysis Center of Shanghai Jiao Tong University. UPLC was performed on a ZIC-HILIC column (100 mm × 2.1 mm i.d., 3.5 µm; Merck). The column was eluted with gradient solvent from A: B (5: 95) to A: B (40: 60) at a flow rate of 0.40 mL/min, where A is 50 mM ammonium formate and B is acetonitrile. We employed the following MS experimental parameters: a negative polarity with 2.0 kV capillary voltage, 20 V sampling cone and 6 eV collision energy. c-di-GMP was detected in M-1 ion of *m/z* 689.086 with a fragment ion of *m/z* 344.040, and cGMP as an input in M-1 ion of *m/z* 344.039.

### The SILAC MS assay of quantitative acetylation proteomics

*E.coli* BW25113 (*△lysA*) and BW25113 with YdeH overexpression (*△lysA ydeH*^+^) were subjected for SILAC MS assay. The two strains were activated in LB medium, and then were cultured in 2% glucose M9 minimal media supplemented with heavy isotopes of lysine (^13^C_6_^14^N_2_-lysine) or light isotopes of lysine (^12^C_6_^14^N_2_-lysine) (Silantes, Munich, Germany) for *△lysA* and *△lysA ydeH*^+^, respectively. Strains were induced by 0.1 mM IPTG during exponential growth (OD 600nm = ~0.4) and the cells were harvested after inducing for 4 h (OD 600nm = ~1.0). These cells (~30 OD) were added with 1 mL lysis buffer (8 M Urea, 100 mM NH_4_HCO_3_, 2 mM sodium butyrate, 5 mM nicotinamide, 1x protease inhibitor (Roche, Basel, Switzerland), pH 8.0) and lysed for 2 min at 4°C by an Ultrasonic Cell Disruptor (Cheng-cheng Weiye Science and Technology, Beijing, China). Protein concentration was determined by BCA kit (Pierce, MA, USA). The labeling efficiency of *E.coli* cultured in “heavy” medium was checked before sequential proteomic experiments.

Light-labeled and heavy-labeled lysate were equally mixed. Cysteine bonds were reduced by 5 mM dithiothreitol (DTT) at 56°C for 30 min and followed by alkylation reaction with 15 mM iodoacetamide at room temperature in darkness for 30 min. The alkylation reaction was quenched by 30 mM cysteine. The protein solution was diluted to less than 2 M Urea by addition of 100 mM NH_4_HCO_3_ (pH 8.0) and then digested with sequencing grade trypsin at a trypsin-to-protein ratio of 1: 50 (w/w) at 37°C for 16 h. For complete digestion, additional trypsin was added at trypsin-to-protein ratio of 1: 100 (w/w) at 37°C for another four hours. The tryptic peptides were desalted through SepPak C18 cartridges (Waters, MA, USA) and vacuum dried.

To enrich the lysine acetylated peptides, 2 mg desalted peptides were dissolved in NETN buffer (600 mM NaCl, 1 mM EDTA, 50 mM Tris-HCl, 0.5% NP-40, pH 8.0) and incubated with 10 μL drained pre-washed anti-acetyl beads (Immunechem, Burnaby, Canada) at 4°C overnight with gentle shaking. The beads were gently washed for four times with NETN buffer and twice with deionized water. The bound peptides were eluted with 0.1% TFA and vacuum dried. The eluted peptides were desalted with C18 ZipTips (Millipore, MA, USA) according to the manufacturer’s instructions.

Enriched acetylated peptides were analyzed by nano flow LC-MS/MS using an EASY-nLC 1000 system connected to Orbitrap Fusion mass spectrometer (Thermo Fisher Scientific, MA, USA). Peptides were dissolved in solvent A (0.1% FA in 2% ACN) and separated using a homemade reverse-phase C18 analytical column (75 μm ID × 18 cm length, 3 μm particle size) with a 90-min gradient from 5% to 80% solvent B (0.1% FA in 90% ACN) at a constant flow rate of 300 nL/min. Intact peptides with m/z 300-1400 were detected at a resolution of 120,000 at m/z 200. Ions with intensity above 5,000 were isolated and sequentially fragmentized by Higher Collision Dissociation (HCD) with normalized collision energy of 32% in top speed mode. The automatic gain control (AGC) targets were set at 5.0e5 for full scan and 7.0e3 for MS/MS scan, respectively. The dynamic exclusion duration was set as 30s.

MS/MS data files were processed with MaxQuant software (version 1.5.3.8) against *Escherichia coli* (strain K12) database from Uniprot (proteome ID: UP000000625, 4309 sequences, last modified on May 13th, 2017) with a reversed decoy database. SILAC was selected as “doublets” and “Heavy labels” panel was selected as heavy lysine (Lys6). Trypsin/P was chosen as the digestion enzyme and two maximum missing cleavages was allowed. Carbamidomethyl (C) was specified as the fixed modification and variable modifications were oxidation (M), acetylation (Protein N-term) and acetylation (K). False discovery rates (FDR) at protein, peptide and modification level were all set as 1%. For quantitative analysis, the normalized H/L ratio of each acetylated peptide exported by MaxQuant software was corrected at the protein level to eliminate the protein abundance difference. The SILAC mass spectrometry proteomics raw data have been deposited to the ProteomeXchange^63^ Consortium via the PRIDE^64^ partner repository with the dataset identifier PXD007616 (Username, Password: dD4exhSO).

### Identification of deacetylation sites by Q Exactive plus MS

Five proteins of the DGC pathway, YdeH, Era, YegE, YeaJ and YihK were constructed to Pca24N and purified from *E.coli* BL21. After SDS-PAGE separation and tryptic digestion, these proteins were mixed and analyzed by mass spectrometry. Briefly, nanoLC−MS/MS-experiments were performed on an EASY-nLC system (Thermo Scientific, Odense, Denmark) connected to a Q Exactive Plus (Thermo Scientific, Bremen, Germany) through a nanoelectrospray ion source. Samples (1 µL) were loaded by an autosampler onto a 2-cm packed pre-column (75 µm ID x 360 µm OD) in 0.1% HCOOH/water (buffer A) at a flow rate of 1 µL/min for 5 min. Analytical separation was performed over a 15-cm packed column (75 µm ID x 360 µm OD) at 300 nL/min with a 60 mins gradient of increasing CH3CN (buffer B, 0.1% HCOOH/CH_3_CN). Both pre-column (5 µm diameter, 200 Å pore size) and analytical column (3 µm diameter, 100 Å pore size) were packed with C18-reversed phase silica (DIKMA-inspire TM, CA, USA) using a pressure bomb. Following sample loading, buffer B was increased rapidly from 3% to 6% over 5min and then shallowly to 22% over 36 min, and then to 35% over 9 min followed by a quick increase to 95% over 3min, and hold at 95% for 7 min. The total acquisition duration lasted for 60 min. The Q Exactive Plus mass spectrometer was operated in the data dependent mode to automatically switch between full scan MS and MS/MS acquisition. Survey full scan MS spectra (m/z 350−1800) were acquired in the Orbitrap with 70 000 resolution (m/z 200) after accumulation of ions to a 3×10 6 target value based on predictive AGC from the previous full scan. Dynamic exclusion was set to 60 s. The 15 most intense multiply charged ions (z≥2) were sequentially isolated and fragmented in the octupole collision cell by higher-energy collisional dissociation (HCD) with affixed injection time of 55 ms and 17500 resolutions for the fast scanning method. Typical mass spectrometric conditions were as follows: spray voltage, 1.7 kV; heated capillary temperature, 320°C; normalized HCD collision energy 27%. The MS/MS ion selection threshold was set to 9×10^3^ counts. A 1.6 Da isolation width for the samp7les was chosen. The mass spectrometry raw data have been submitted to PRIDE^63, 64^ with project accession PXD007651 (User, password: 6rxEMkjm). And the raw data file was named “20161202_ZHN_AC.raw” with the search file “20161202_ZHN_AC-01.msf”.

YdeH was chosen for in-depth quantitative MS analysis using Q Exactive plus mass spectrometer. YdeH was overexpressed in *E. coli* and affinity purified. YdeH was treated with CobB for deacetylation and untreated WT YdeH as the control. After SDS-PAGE separation and trypsin digestion, the samples were analyzed by Q Exactive plus mass spectrometer under the same experiment condition. The MS raw data was processed using Protein Discovery software (ThermoFisher) to identify the lysine acetylation sites. The acetylation sites, which were identified in untreated WT YdeH sample but not in CobB treated sample, were considered to be deacetylated by CobB. The mass spectrometry raw data have been submitted to PRIDE with project accession PXD008113 (Username, password: jKBngFLH).

### Determination of the deacetylation activity of CobB *in vivo*

The strains, △ *cobB*, WT, *ydeH*^+^, *ydeH*^G206A,G207A^, △ *cobB::cobB*, △*cobB::cobB ydeH*^+^ were used for to assay the deacetylation activity of CobB. These strains were grown in Vogel-Bonner medium (0.81 mM MgSO_4_·7H_2_O, 43.8 mM K_2_HPO_4_, 10 mM C_6_H_8_O_7_·H_2_O, 16.7 mM NaNH_4_HPO_4_·4H_2_O) with 10 mM acetate at 25°C for 12 h and induced by 0.2 mM IPTG at 25°C for 12 h. 20 OD cells were harvested and repeatedly freeze-thawed three times. The cells were then treated with 4mL lysis buffer (50 mM NaH_2_PO_4_, 300 mM NaCl, 10 mM imidazole (Sigma-Aldrich), 1mg/mL lysozyme (Sangon Biotech, Shanghai, China), 1x CelLytic B (Sigma-Aldrich), 50 units/mL of Benzonase (Sigma-Aldrich) and 1 mM PMSF (Sigma-Aldrich), pH 8.0) at 4°C for 20 min with vigorous shaking. After lysis and centrifugation at 10,000 rpm for 5 min, 20 μL anti-FLAG antibody (Sigma-Aldrich) and 50 μL protein G conjugated agarose beads (Roche) were added at 4°C for 2 h with gentle agitation to enrich the 3xFLAG-tagged Acs. Protein G conjugated agarose was harvested and washed three times by buffer A (50 mM NaH_2_PO_4_, 300 mM NaCl, 10 mM imidazole, pH 8.0). The 3xFLAG-tagged Acs was then eluted by heating at 95°C for 10 min. The amount of Acs protein and deacetylation level of these samples were analyzed by western blotting. Membranes were incubated with the pan anti-acetyl antibody (Cell Signaling Technology with a 1:1000 dilution) for at 4°C for 16 h and an anti-FLAG antibody (Sigma-Aldrich with a 1:2000 dilution) at 4°C for 16 h. The IRDye 800 antibody was used as a secondary antibody, and membranes were incubated for 1 h at room temperature. The membranes were washed three times in TBST between each antibody incubation step. Final visualization was performed using an Odyssey Infrared Imaging System.

### Determination of the strain growth curve in Vogel-Bonner medium

The strains described above were grown in Vogel-Bonner medium with 10 or 30 mM acetate or propionate at 25°C for 12 h and then induced by 0.2 mM IPTG at 25°C for 20 h. During the entire 32 h growth period, the cell concentrations were measured at OD_600_ using Nanodrop 2000s at 8, 12, 16, 24 and 32 h. The growth curves were then drawn using GraphPad Prism 6.

### Homology analysis, phylogenetic tree construction, and sequence alignment

The 25 bacteria that we selected for homology analysis were *Escherichia coli, Salmonella typhimurium, Yersinia pestis, Shigella dysenteriae, Citrobacter amalonaticus, Erwinia tracheiphila, Vibrio parahaemolyticus, Klebsiella oxytoca, Cedecea neteri, Achromobacter sp.ATCC35328, Siccibacter colletis, Lelliottia amnigena, Buttiauxella brennerae, Kluyvera ascorbata, Mangrovibacter sp.MFB070, Cronobacter dublinensis, Hafnia alvei, Chania multitudinisentens, Serratia symbiotica, Yokenella regensburgei, Trabulsiella odontotermitis, Leclercia adecarboxylata, Pantoea dispersa, Shimwellia blattae, Kosakonia radicincitans*. The CobB sequences were examined at the NCBI website and the phylogenetic tree was constructed by EMBL-EBI Clustalw2 online tool. Sequence alignment was performed using DNAMAN 2.0.

### Determination of the YdeH DGC activity

The purified WT YdeH 5 uM (0.14mg/ml) protein was incubated with CobB 1.6 uM (0.05mg/ml) for deacetylating at 37℃ for 30 min. For DGC activity determination, the WT YdeH, deacetylated YdeH and YdeH mutants were incubated with 1 mM GTP, 5 mM MgCl_2_ in 100 ul at 30℃ for 2 h. The reaction was terminated with heating at 95℃ for 10 min and spun for 10 min at 18,000 g to separate the enzyme from the reactions. Before analyzed, we add 2 uM cGMP to these reactions as the control. The c-di-GMP concentrations were determined by UPLC-IM-MS with the same parameters as mentioned earlier.

### Determination of the YdeH K4 acetylation stoichiometry using AQUA quantification

We used the absolute quantification (AQUA) method^39, 40^ to quantify the peptide levels. We synthesized the AQUA peptides for deacetylated (KTTEIDAI**L**(^13^C_6_,^15^N)LNLNK and TTEIDAI**L**(^13^C_6_,^15^N)LNLNK) and acetylated peptides (K^ac^TTEIDAI**L**(^13^C_6_,^15^N)LNLNK) of YdeH K4. To acquire the endogenous YdeH protein, we inserted a chromosomal C-terminal Flag-tag to YdeH and immuno-precipitated by an anti-FLAG antibody. The strains, △*cobB* and WT, were used for to assay the YdeH K4 acetylation stoichiometry. These strains were grown in LB medium overnight and transfer to 1 L VB-E medium with 1:500 dilution and grown to OD600 = ~0.3 at 25℃ for 16 h as the exponential growth cells and to OD600 = ~0.6 at 25℃ for 42 h as the stationary phase cells. These cells were harvested and lyzed by high pressure with 50 mL lysis buffer (50 mM NaH_2_PO_4_, 300 mM NaCl, and 1 mM PMSF (Sigma-Aldrich), pH 8.0). After lysis and centrifugation at 10,000 rpm for 5 min, 100 μL anti-FLAG antibody (Sigma-Aldrich) was added and incubated at 4°C for 20 h with gentle agitation to enrich the 3xFLAG-tagged YdeH. 300 μL protein G conjugated agarose beads (Roche) were added and incubated at 4°C for 4 h with gentle agitation to enrich the anti-Flag antibody. Protein G conjugated agarose was harvested and washed for three times by buffer A (50 mM NaH_2_PO_4_, 300 mM NaCl, pH 8.0). The 3xFLAG-tagged YdeH was eluted by 2 mL 0.2 mg/mL Flag peptide at 4°C for 4 h. The free Flag peptide was separated through dialysis.

To acquire the overexpressed YdeH, we constructed the YdeH to pCA24N in △*cobB* strain (△*cobB ydeH^+^*). This strain was grown in LB medium overnight and transfer to 100 mL LB medium with 1:100 dilution and grown to OD600 = ~0.6 at 37℃ for 2 h, 0.1 mM IPTG was added to induce at 37℃ for 2 h. The cell was harvested and lyzed with 35 mL lysis buffer. After lysis and centrifugation at 10,000 rpm for 5 min, 1 mL Ni-IDA beads (Senhuimicrosphere, Suzhou, China) was added and incubated at 4°C for 1 h with gentle agitation to enrich the His-tagged YdeH. The His-tagged YdeH was eluted by 250 mM imidazole at 4°C for 20 min. The free imidazole was separated through dialysis.

For YdeH acetylation stoichiometry, affinity purified YdeH proteins were digested by trypsin overnight after cysteine reduction and alkylation reaction. Then tryptic peptides were desalted. The synthetic AQUA standard peptides were spiked into the digested YdeH samples with the close MS intensity to the native peptides. Peptide mixture containing AQUA peptides were analyzed by Q-Exactive mass spectrometer with three independent measurements. Full MS scan mode was used and the scan range was set as 700 to 900 m/z, containing all three targeted peptides. Extracted ion chromatography (XIC) peak areas of native peptide and corresponding heavy labeled peptide were used for stoichiometry calculation. The mass spectrometry proteomics raw data have been deposited to the ProteomeXchange^63^ Consortium via the PRIDE^64^ partner repository with the dataset identifier PXD007616 (Username, Password: dD4exhSO).

### Determination of stability of YdeH *in vitro*

After overexpressed in CobB deficient cells, YdeH was purified and diluted to 0.14 mg/mL, followed with or without CobB treatment. These samples were incubated at 37℃ for deacetylation for 0.5 h. Then 1 mM GTP was added, the samples were incubated at 30℃ for another 1 h and 2h. These samples were centrifuged at 12,000 g for 10 min to separate the soluble protein from the precipitation.

For YdeH mutants, these were diluted to 0.14 mg/mL incubated at 37℃ for 0.5 h. Then 1 mM GTP was added, the samples were incubated at 30℃ for another 1 h and 2h. These samples were centrifuged at 12,000 g for 10 min to separate the soluble protein from the precipitation.

### Statistical analysis

Graphs were plotted using GraphPad Prism 6 and the statistical analyses were performed using Excel. Pairwise comparisons were performed using two-tailed Student’s *t*-test, and statistical significance was set at *P* <0.05. Error bars represent the mean ± standard errors of mean (S.E.M.).

## Acknowledgements

We thank Prof. Yufeng Yao of Shanghai Jiao Tong University for providing *S. typhimurium* CobB overexpression plasmid. We thank Prof. Yan Wei and Liyun Ji of Shanghai Jiao Tong University for performing the Q Exactive plus assay. We thank Ademi Zhakyp of Nazarbayev University for proof-reading the manuscript. This study was supported in part by The National Key Research and Development Program of China Grant 2016YFA0500600, National Natural Science Foundation of China Grants 31670831, 31370813, to S.C.T., 31370750, 31670722 to D.M.C., and 31670066 to M.J.T., and National Basic Research Program of China (973 Program) (No. 2014CBA02004) to M.J.T..

## Author Contributions Statement

S.C.T. conceived the idea. L.J.B. provided key reagents. Z.W.X. and C.X.L. performed interaction assay and functional analysis. Z.W.X. and C.X.L. performed enzyme activity assay. Z.W.X., H.N.Z., X.R.Z. and H.T.L. performed structure analysis and protein mutation. H.N.Z. and X.R.Z. performed the ITC and MST assay. Z.W.X., H.N.Z., X.R.Z., H.W.J., S.J.G and F.L.W. prepared the figures with the help of S.H.W. L.L.Q. and M.J.T. performed the SILAC-MS. F.L. performed the UPLC-IM-MS analysis. J.L.H. performed Nano LC maXis impact UHR-TOF MS analysis. Z.W.X., H.N.Z., X.R.Z., D.M.C. and S.C.T. wrote the manuscript.

## References

1. ROSS, P. et al. Regulation of cellulose synthesis in Acetobacter xylinum by cyclic diguanylic acid. Nature 325, 279–281 (1987).

2. Hengge, R. Principles of c-di-GMP signalling in bacteria. Nature reviews. Microbiology 7, 263–273 (2009).

3. Jenal, U., Reinders, A. & Lori, C. Cyclic di-GMP: second messenger extraordinaire. Nature reviews. Microbiology (2017).

4. Bush, M.J., Tschowri, N., Schlimpert, S., Flardh, K. & Buttner, M.J. c-di-GMP signalling and the regulation of developmental transitions in streptomycetes. Nature reviews. Microbiology 13, 749–760 (2015).

5. Nesper, J., Reinders, A., Glatter, T., Schmidt, A. & Jenal, U. A novel capture compound for the identification and analysis of cyclic di-GMP binding proteins. Journal of Proteomics 75, 4874–4878 (2012).

6. Paul, K., Nieto, V., Carlquist, W.C., Blair, D.F. & Harshey, R.M. The c-di-GMP binding protein YcgR controls flagellar motor direction and speed to affect chemotaxis by a "backstop brake" mechanism. Molecular cell 38, 128–139 (2010).

7. Lori, C. et al. Cyclic di-GMP acts as a cell cycle oscillator to drive chromosome replication. Nature 523, 236–239 (2015).

8. Tschowri, N. et al. Tetrameric c-di-GMP mediates effective transcription factor dimerization to control Streptomyces development. Cell 158, 1136–1147 (2014).

9. Nesper, J. et al. Cyclic di-GMP differentially tunes a bacterial flagellar motor through a novel class of CheY-like regulators. Elife 6 (2017).

10. Schirmer, T. C-di-GMP Synthesis: Structural Aspects of Evolution, Catalysis and Regulation. J. Mol. Biol. 428, 3683–3701 (2016).

11. Ryan, R.P. et al. Cell-cell signaling in Xanthomonas campestris involves an HD-GYP domain protein that functions in cyclic di-GMP turnover. Proceedings of the National Academy of Sciences of the United States of America 103, 6712–6717 (2006).

12. Schmidt, A.J., Ryjenkov, D.A. & Gomelsky, M. The ubiquitous protein domain EAL is a cyclic diguanylate-specific phosphodiesterase: Enzymatically active and inactive EAL domains. Journal of bacteriology 187, 4774–4781 (2005).

13. Christen, M., Christen, B., Folcher, M., Schauerte, A. & Jenal, U. Identification and characterization of a cyclic di-GMP-specific phosphodiesterase and its allosteric control by GTP. J. Biol. Chem. 280, 30829–30837 (2005).

14. Boehm, A. et al. Second messenger signalling governs Escherichia coli biofilm induction upon ribosomal stress. Molecular microbiology 72, 1500–1516 (2009).

15. Tuckerman, J.R. et al. An oxygen-sensing diguanylate cyclase and phosphodiesterase couple for c-di-GMP control. Biochemistry 48, 9764–9774 (2009).

16. Ko, M. & Park, C. Two novel flagellar components and H-NS are involved in the motor function of Escherichia coli. J Mol Biol 303, 371–382 (2000).

17. Doerks, T., Copley, R.R., Schultz, J., Ponting, C.P. & Bork, P. Systematic identification of novel protein domain families associated with nuclear functions. Genome research 12, 47–56 (2002).

18. Spangler, C., Kaever, V. & Seifert, R. Interaction of the diguanylate cyclase YdeH of Escherichia coli with 2’,(3’)-substituted purine and pyrimidine nucleotides. The Journal of pharmacology and experimental therapeutics 336, 234–241 (2011).

19. Imai, S. & Guarente, L. Ten years of NAD-dependent SIR2 family deacetylases: implications for metabolic diseases. Trends Pharmacol. Sci. 31, 212–220 (2010).

20. Greiss, S. & Gartner, A. Sirtuin/Sir2 phylogeny, evolutionary considerations and structural conservation. Mol. Cells 28, 407–415 (2009).

21. Tucker, A.C. & Escalante-Semerena, J.C. Biologically Active Isoforms of CobB Sirtuin Deacetylase in Salmonella enterica and Erwinia amylovora. Journal of bacteriology 192, 6200–6208 (2010).

22. Starai, V.J., Celic, I., Cole, R.N., Boeke, J.D. & Escalante-Semerena, J.C. Sir2-dependent activation of acetyl-CoA synthetase by deacetylation of active lysine. Science 298, 2390–2392 (2002).

23. Li, R. et al. CobB regulates Escherichia coli chemotaxis by deacetylating the response regulator CheY. Molecular microbiology 76, 1162–1174 (2010).

24. Zhang, Q.F. et al. Reversibly acetylated lysine residues play important roles in the enzymatic activity of Escherichia coli N-hydroxyarylamine O-acetyltransferase. The FEBS journal 280, 1966–1979 (2013).

25. Zhou, Q., Zhou, Y.N., Jin, D.J. & Tse-Dinh, Y.-C. Deacetylation of topoisomerase I is an important physiological function of E. coli CobB. Nucleic acids research (2017).

26. Hentchel, K.L. & Escalante-Semerena, J.C. Complex regulation of the sirtuin-dependent reversible lysine acetylation system of Salmonella enterica. Microbial cell (Graz, Austria) 2, 451–453 (2015).

27. Hentchel, K.L., Thao, S., Intile, P.J. & Escalante-Semerena, J.C. Deciphering the Regulatory Circuitry That Controls Reversible Lysine Acetylation in Salmonella enterica. Mbio 6 (2015).

28. Avalos, J.L., Bever, K.M. & Wolberger, C. Mechanism of sirtuin inhibition by nicotinamide: altering the NAD(+) cosubstrate specificity of a Sir2 enzyme. Molecular cell 17, 855–868 (2005).

29. Chen, C.S. et al. A proteome chip approach reveals new DNA damage recognition activities in Escherichia coli. Nature methods 5, 69–74 (2008).

30. Kramer, K. et al. Photo-cross-linking and high-resolution mass spectrometry for assignment of RNA-binding sites in RNA-binding proteins. Nature methods 11, 1064–1070 (2014).

31. Shu, C., Yi, G., Watts, T., Kao, C.C. & Li, P. Structure of STING bound to cyclic di-GMP reveals the mechanism of cyclic dinucleotide recognition by the immune system. Nature structural & molecular biology 19, 722–724 (2012).

32. Tu, S. et al. YcgC represents a new protein deacetylase family in prokaryotes. Elife 4 (2015).

33. Liu, F.Y., Gu, J., Wang, X.D., Zhang, X.E. & Deng, J.Y. Acs is essential for propionate utilization in Escherichia coli. Biochem. Biophys. Res. Commun. 449, 272–277 (2014).

34. Weinert, B.T. et al. Acetyl-phosphate is a critical determinant of lysine acetylation in E. coli. Molecular cell 51, 265–272 (2013).

35. Castano-Cerezo, S. et al. Protein acetylation affects acetate metabolism, motility and acid stress response in Escherichia coli. Molecular systems biology 10, 762 (2014).

36. Chou, S.H. & Galperin, M.Y. Diversity of Cyclic Di-GMP-Binding Proteins and Mechanisms. Journal of bacteriology 198, 32–46 (2016).

37. Wang, Q.J. et al. Acetylation of Metabolic Enzymes Coordinates Carbon Source Utilization and Metabolic Flux. Science 327, 1004–1007 (2010).

38. Weinert, B.T. et al. Accurate quantification of site-specific acetylation stoichiometry reveals the impact of sirtuin deacetylase CobB on the E. coli acetylome. Molecular & cellular proteomics : MCP (2017).

39. Gerber, S.A., Rush, J., Stemman, O., Kirschner, M.W. & Gygi, S.P. Absolute quantification of proteins and phosphoproteins from cell lysates by tandem MS. Proceedings of the National Academy of Sciences of the United States of America 100, 6940–6945 (2003).

40. Weinert, B.T., Moustafa, T., Iesmantavicius, V., Zechner, R. & Choudhary, C. Analysis of acetylation stoichiometry suggests that SIRT3 repairs nonenzymatic acetylation lesions. The EMBO journal 34, 2620–2632 (2015).

41. Qian, M.X. et al. Acetylation-mediated proteasomal degradation of core histones during DNA repair and spermatogenesis. Cell 153, 1012–1024 (2013).

42. Cohen, T.J. et al. The acetylation of tau inhibits its function and promotes pathological tau aggregation. Nat Commun 2, 252 (2011).

43. Cohen, T.J. et al. An acetylation switch controls TDP-43 function and aggregation propensity. Nat Commun 6, 5845 (2015).

44. Barends, T.R. et al. Structure and mechanism of a bacterial light-regulated cyclic nucleotide phosphodiesterase. Nature 459, 1015–1018 (2009).

45. Zhang, H.-N. et al. Cyclic di-GMP regulates Mycobacterium tuberculosis resistance to ethionamide. Scientific Reports 7 (2017).

46. Dubey, B.N. et al. Cyclic di-GMP mediates a histidine kinase/phosphatase switch by noncovalent domain cross-linking. Sci. Adv. 2 (2016).

47. Christen, M. et al. Asymmetrical distribution of the second messenger c-di-GMP upon bacterial cell division. Science 328, 1295–1297 (2010).

48. Zimmerman, S.B. & Trach, S.O. Estimation of macromolecule concentrations and excluded volume effects for the cytoplasm of Escherichia coli. J. Mol. Biol. 222, 599–620 (1991).

49. Parry, Bra.ley R. et al. The Bacterial Cytoplasm Has Glass-like Properties and Is Fluidized by Metabolic Activity. Cell 156, 183–194 (2014).

50. Kim, G.-W. & Yang, X.-J. Comprehensive lysine acetylomes emerging from bacteria to humans. Trends in Biochemical Sciences 36, 211–218 (2011).

51. Sasikaran, J., Ziemski, M., Zadora, P.K., Fleig, A. & Berg, I.A. Bacterial itaconate degradation promotes pathogenicity. Nature Chemical Biology 10, 371–U382 (2014).

52. Liang, W.X., Malhotra, A. & Deutscher, M.P. Acetylation Regulates the Stability of a Bacterial Protein: Growth Stage-Dependent Modification of RNase R. Molecular cell 44, 160–166 (2011).

53. Caimano, M.J., Drecktrah, D., Kung, F. & Samuels, D.S. Interaction of the Lyme disease spirochete with its tick vector. Cellular microbiology 18, 919–927 (2016).

54. Rotem, O. et al. An Extended Cyclic Di-GMP Network in the Predatory Bacterium Bdellovibrio bacteriovorus. Journal of bacteriology 198, 127–137 (2016).

55. Jonas, K. et al. The RNA binding protein CsrA controls cyclic di-GMP metabolism by directly regulating the expression of GGDEF proteins. Molecular microbiology 70, 236–257 (2008).

56. Zahringer, F., Lacanna, E., Jenal, U., Schirmer, T. & Boehm, A. Structure and signaling mechanism of a zinc-sensory diguanylate cyclase. Structure 21, 1149–1157 (2013).

57. Thauer, R.K., Jungermann, K. & Decker, K. Energy conservation in chemotrophic anaerobic bacteria. Bacteriological reviews 41, 100–180 (1977).

58. Dubbs, J.M. & Robert Tabita, F. Regulators of nonsulfur purple phototrophic bacteria and the interactive control of CO2 as similation, nitrogen fixation, hydrogen metabolism and energy generation. FEMS microbiology reviews 28, 353–376 (2004).

59. Schwer, B. & Verdin, E. Conserved metabolic regulatory functions of sirtuins. Cell Metab. 7, 104–112 (2008).

60. Gentner, M., Allan, M.G., Zaehringer, F., Schirmer, T. & Grzesiek, S. Oligomer formation of the bacterial second messenger c-di-GMP: reaction rates and equilibrium constants indicate a monomeric state at physiological concentrations. Journal of the American Chemical Society 134, 1019–1029 (2012).

61. Poteete, A.R. What makes the bacteriophage lambda Red system useful for genetic engineering: molecular mechanism and biological function. FEMS Microbiol. Lett. 201, 9–14 (2001).

62. Spangler, C., Bohm, A., Jenal, U., Seifert, R. & Kaever, V. A liquid chromatography-coupled tandem mass spectrometry method for quantitation of cyclic di-guanosine monophosphate. Journal of microbiological methods 81, 226–231 (2010).

63. Deutsch, E.W. et al. The ProteomeXchange consortium in 2017: supporting the cultural change in proteomics public data deposition. Nucleic acids research 45, D1100–D1106 (2017).

64. Vizcaino, J.A. et al. 2016 update of the PRIDE database and its related tools. Nucleic acids research 44, D447–D456 (2016).

